# Post-prediction Inference

**DOI:** 10.1101/2020.01.21.914002

**Authors:** Siruo Wang, Tyler H. McCormick, Jeffrey T. Leek

## Abstract

Many modern problems in medicine and public health leverage machine learning methods to predict outcomes based on observable covariates [1, 2, 3, 4]. In an increasingly wide array of settings, these predicted outcomes are used in subsequent statistical analysis, often without accounting for the distinction between observed and predicted outcomes [1, 5, 6, 7, 8, 9]. We call inference with predicted outcomes *post-prediction inference*. In this paper, we develop methods for correcting statistical inference using outcomes predicted with an arbitrary machine learning method. Rather than trying to derive the correction from the first principles for each machine learning tool, we make the observation that there is typically a low-dimensional and easily modeled representation of the relationship between the observed and predicted outcomes. We build an approach for the *post-prediction inference* that naturally fits into the standard machine learning framework where the data is divided into training, testing, and validation sets. We train the prediction model in the training set, estimate the relationship between the observed and predicted outcomes on the testing set, and use that model of the relationship to correct inference on the validation set and subsequent statistical models. We show our *postpi* approach can correct bias and improve variance estimation (and thus subsequent statistical inference) with predicted outcome data. To show the broad range of applicability of our approach, we show *postpi* can improve inference in two totally distinct fields: modeling predicted phenotypes in re-purposed gene expression data [10] and modeling predicted causes of death in verbal autopsy data [11]. We have made our method available through an open-source R package: [https://github.com/leekgroup/postpi]

## 1 Significance Statement

Machine learning is now being used across the entire scientific enterprise. Researchers commonly use the predictions from these models in downstream statistical analysis as if they were observed data. We show that this can lead to extreme bias and uncontrolled variance in downstream statistical models. We propose a new statistical adjustment to correct biased inference in regression models using predicted outcomes – regardless of the machine learning model used to make those predictions.

## 2 Introduction

The past decade has seen both an explosion in data available for precision health [12, 13, 14] and, simultaneously, user-friendly tools such as the caret package [15] and Scikit-learn [16] that make implementing complex statistical and machine learning methods possible for an increasingly wide range of scientists. For example, machine learning from electronic medical records is used to predict phenotypes [1, 17], genomic data is used to predict health outcomes [2], survey data is used to predict cause of death in settings where deaths happen outside of hospitals [3, 18]. The increased focus on ideas like precision medicine means the role of machine learning in medicine and public health will only increase [4]. As machine learning plays an increasingly critical role across scientific disciplines, it is critical to consider all sources of potential variability in downstream inference to ensure stable statistical results [19].

In many settings, researchers do not observe outcomes directly, so observed outcomes are often replaced with predicted outcomes from machine learning models in downstream analyses [1, 5, 6, 7, 8, 9]. One example from genetics is association studies between genetic variants and Alzheimer’s disease for young adults. Because young adults have not developed Alzheimer’s disease, it is difficult to associate the phenotype with genetic variants. However, these adults’ older relatives can be used to predict the ultimate phenotype of participants in the study using known inheritance patterns for the disease. The predicted outcome can be used in place of the observed Alzheimer’s status when performing a genome-wide association study [6].

This is just one example of the phenomenon of *post-prediction inference* or *posti* for short. Though common, this approach poses multiple potential statistical challenges. The predicted outcomes may be biased, may have less variability than the actual outcomes, or the predictions may use the same covariates used as independent variables in subsequent statistical analyses leading to overfitting. Standard practice in many applications is to treat predicted outcomes as if they were the observed outcomes in subsequent regression models [1, 5, 6, 7, 8, 9]. As we will show, uncorrected post-prediction inference will frequently have deflated standard errors, bias, and inflated false positive rates.

Post-prediction inference appears across fields and has been recognized as a potential source of error in recent work on prevalence estimation (see for example [20] and [21] in the context of data set shift and [22] in document class prevalence estimation). Here, we focus on developing analytical and bootstrap-based approaches to correct regression estimates, standard errors, and test statistics in inferential regression models using predicted outcomes. We focus specifically on settings where a predicted outcome becomes the dependent variable in the subsequent inferential regression analysis. We derive an analytical correction in the case of linear regression, and bootstrap-based corrections for more general regression models, focusing on linear and logistic regression as they are the most common inferential models. Our bootstrap-based approach can, however, easily be extended to any generalized linear regression inference model.

Both our analytical and bootstrap-based corrections take advantage of the standard structure for machine learning problems. We assume that we have at least three separate sub-samples, which we here label as the training set, the testing set, and the validation set (see Figure 1). Our assumption is that the data generating distribution for the three data sets is the same. We assume that the training and testing set are complete – we observe both the outcome of interest (*y*) and the covariates (*x*). In the validation set, we assume that only the covariates are observed. The validation set could represent either a validation subset from a single sample, or a future prospectively collected data set where we wish to perform inference but it is too costly or challenging to collect the outcome.

**Figure 1:**
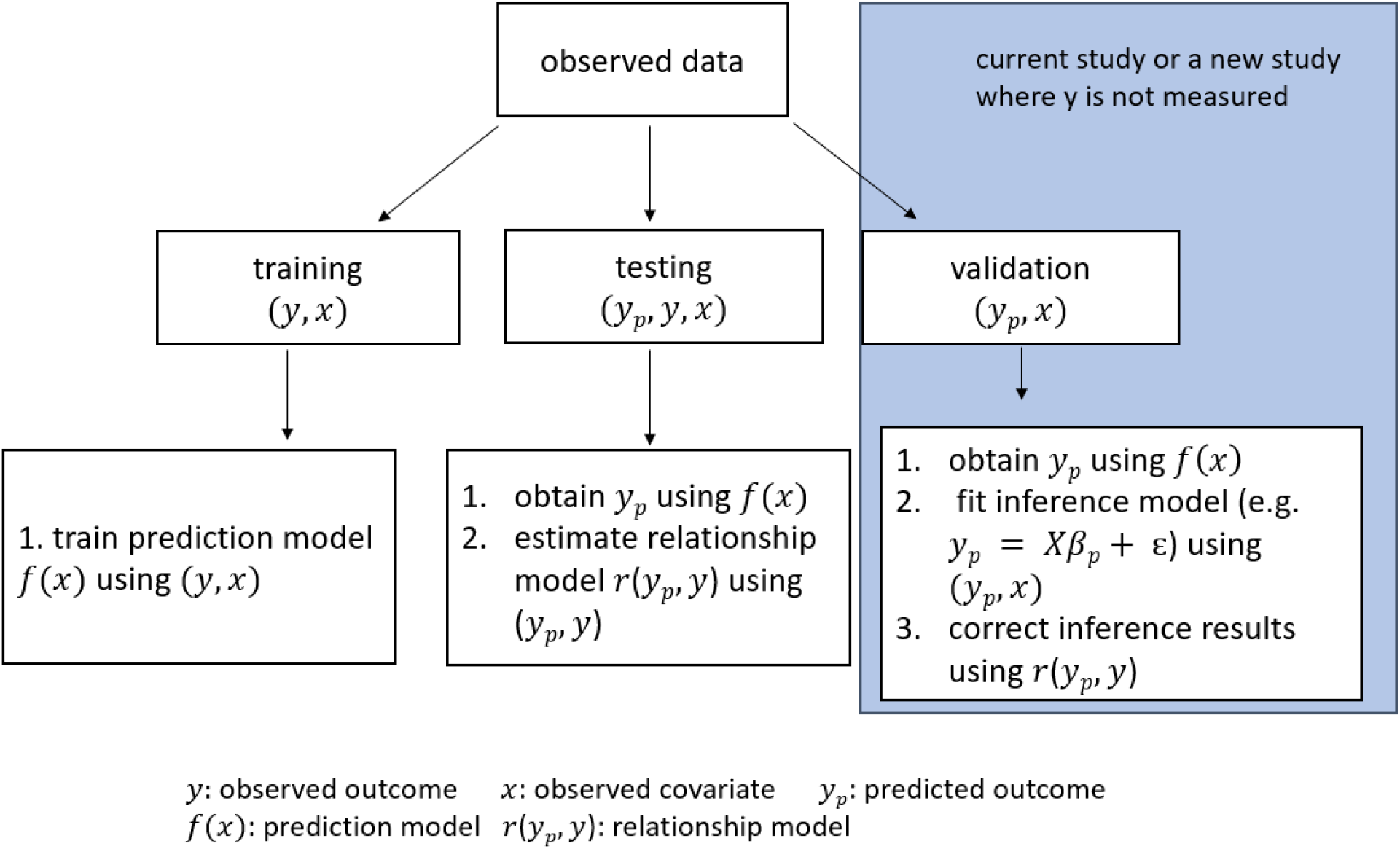
Data split diagram. The common structure of our approach is to divide the observed data into training, testing, and validation sets. The training set is used to train the prediction model, the testing set is used to estimate the relationship between the observed and predicted outcomes, and the validation set is for fitting the downstream inferential model where the relationship model is used to correct inference in the subsequent statistical inference.

A prediction function for the outcome 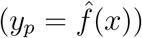 is generated on the training set and applied in the testing and validation sets. In the validation set our goal is to perform inference on a regression model of the form *g*(*E*[*y*|*X*]) = *Xβ*. However, on the validation set, only covariates are observed so instead we must fit *g*(*E*[*y_p_*|*X*]) = *Xβ_p_*. Our goal is to recover the inference we would have obtained if we had observed the true outcomes *y* in the validation set. To correct inference using the predicted outcomes, we take advantage of the testing data set where we have both the predicted (*y_p_*) and observed (*y*) outcomes. We derive a correction for inference using *y_p_* based on the relationship between *y* and *y_p_*.

An advantage of this approach is that it is not specific to a particular machine learning model. That is, we do not need to know *a priori* the expected out of sample performance for a given method. Instead, we assume that the relationship between the predicted and observed outcomes on the testing set well-characterizes the same relationship on the validation set.

The setting we describe has parallels with multiple imputation [23] for missing data, but has several distinct features. Any prediction problem could be cast as a missing data problem where all of the values are missing and no missingness mechanism distinguishes observed and unobserved outcomes. The reason is that on the validation set or the subsequent analyses in practical problems there are no observed outcome data. Multiple imputation also frequently relies on a generative model for simulating data, however in our setting, we wish to build a framework that can be used for any machine learning model, regardless of its operating characteristics. We, therefore, need a new methodology that can use a black-box machine learning algorithm but build a simple model for the relationship between the predicted and observed outcome data. This problem is related to the idea of errors-in-variables [24] or measurement error models [25], where either the outcome or the covariates are measured with error. However, in prediction problems, we can no longer assume that the errors are independent of the predicted values, since the machine learning predictions may be more accurate for subsets of the *y* values.

Aside from its utility in medicine and public health, the methods we propose are also broadly applicable in the social sciences. In political science, for example, machine learning tools classify sentiment or political identification in segments of text and then fit regression models to identify features of text leaning towards one party or another [26]. In urban sociology, researchers used machine learning tools to infer the race of household heads subject to eviction, then used regression models to understand heterogeneity in circumstances related to evictions of individuals of a particular race [27].

We apply our *postpi* approach to two open problems: modeling the relationship between gene expression levels and tissue types [2], and understanding trends in (predicted) cause of death [28, 29]. We show that our method can reduce bias, appropriately model variability, and correct hypothesis testing in the case where only the predicted outcomes are observed. We also discuss the sensitivity of our approach to changes in the study population that might lead to a violation of the assumptions of our approach. Our *postpi* approach is available as an open-source R package available from Github: [https://github.com/leekgroup/postpi].

The remainder of the paper is organized as follows. In the remainder of this section, we provide a simple example of the setting where our approach would be valuable. Then, we present our simulation-based and analytical correction methods in Section 3, followed by evaluation in simulated cases and real applications. We conclude in Section 4.

### 2.1 Illustrative example

We begin with an illustrative simulated example to highlight the issues that can arise with uncorrected post-prediction inference. The methods we present in the subsequent sections cover a wider ranger of settings and do not require the distributional assumptions we make here for exposition. Here we simulate observations for the outcome *y_i_* and covariates *x_ij_* for *i* = 1, …, *n*,*j* = 1, …, *p*. We use *x_i_* to denote vector [*x_i_*_1_, …, *x_ip_*]. In our simulation, we generate data according to the following true relationship between *y* and *x* which we will denote by *f* (·):

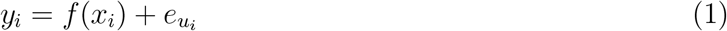

This model represents the true underlying data generating distribution, which is unknown in actual analysis settings.

Linear or generalized linear models are common approaches for perform inference, even when the data generating process is unknown. We use *X_i_* to denote the design matrix in regression models. For example, we may be interested in fitting models of the form:

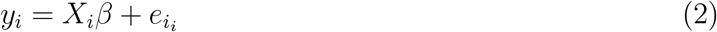

We assume that we are in a data settings where the outcome *y_i_* is too expensive or time consuming to collect. Instead we use a prediction model of the form:

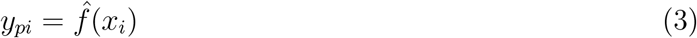

to predict the outcome. The prediction model may be arbitrarily complicated since the goal of the prediction is to minimize a suitable loss function: *E*||*y* − *f* (*x*)||, not to perform inference on the relationship between *y* and *x*.

Then, we fit the regression model of interest, using the predicted outcomes *y_pi_*:

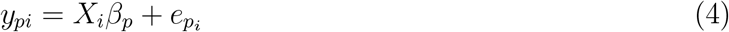

Since the observed outcomes *y_i_* are not available and we, instead, use the predicted *y_pi_* we get a coefficient estimate 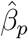 based on the model fit using *y_pi_* as the outcome, such that *E*(*y_pi_*|*X_i_*) = *X_i_β_p_*. Equation (4) no long appropriately reflects our uncertainty about the outcome - leading to to bias in the estimates, standard errors that are too small, and anti-conservatively biased p-values and false positives.

We specify that the state of nature model *f* (·) is a linear function, the prediction model 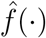 is a random forest [30, 31] machine learning model, and the downstream inferential model is a linear regression model to relate outcomes and covariates. This simulation is designed to reflect standard statistical practice where the data generating distribution is unknown and the target of inference is a regression model. The only difference is that we assume that the outcome is not observed and must be predicted. We do not require these exact distributional assumptions for our methods below, but use this simple case as a means of illustrating the potential issues with post-prediction inference.

Figure 2 shows a simple simulated example of this idea. We simulate covariates *x_i_*_1_, *x_i_*_2_, *x_i_*_3_, *x_i_*_4_ and error terms 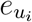 from normal distributions and simulate the observed outcome *y_i_* using the specified state of nature model *f* (·) from a normal distribution with mean function a linear combination of covariates *x_i_*_1_, *x_i_*_2_, *x_i_*_3_, *x_i_*_4_. Then we separate the simulated values into training, testing, and validation sets that have the same data generating systems, and we follow the same procedure in each set as described in the data split diagram in Figure 1. In the training set, we train a random forest [30, 31] model as the prediction model 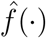 using all covariates *x_i_*_1_, *x_i_*_2_, *x_i_*_3_, *x_i_*_4_ and observed outcome *y_i_*. In the testing set, we apply this prediction model to the observed covariates *x_i_*_1_, *x_i_*_2_, *x_i_*_3_, *x_i_*_4_, in order to obtain the predicted values 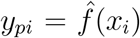. Then we estimate the relationship between the predicted and observed outcomes. In the validation set, we fit a linear regression model as the inference model.

**Figure 2:**
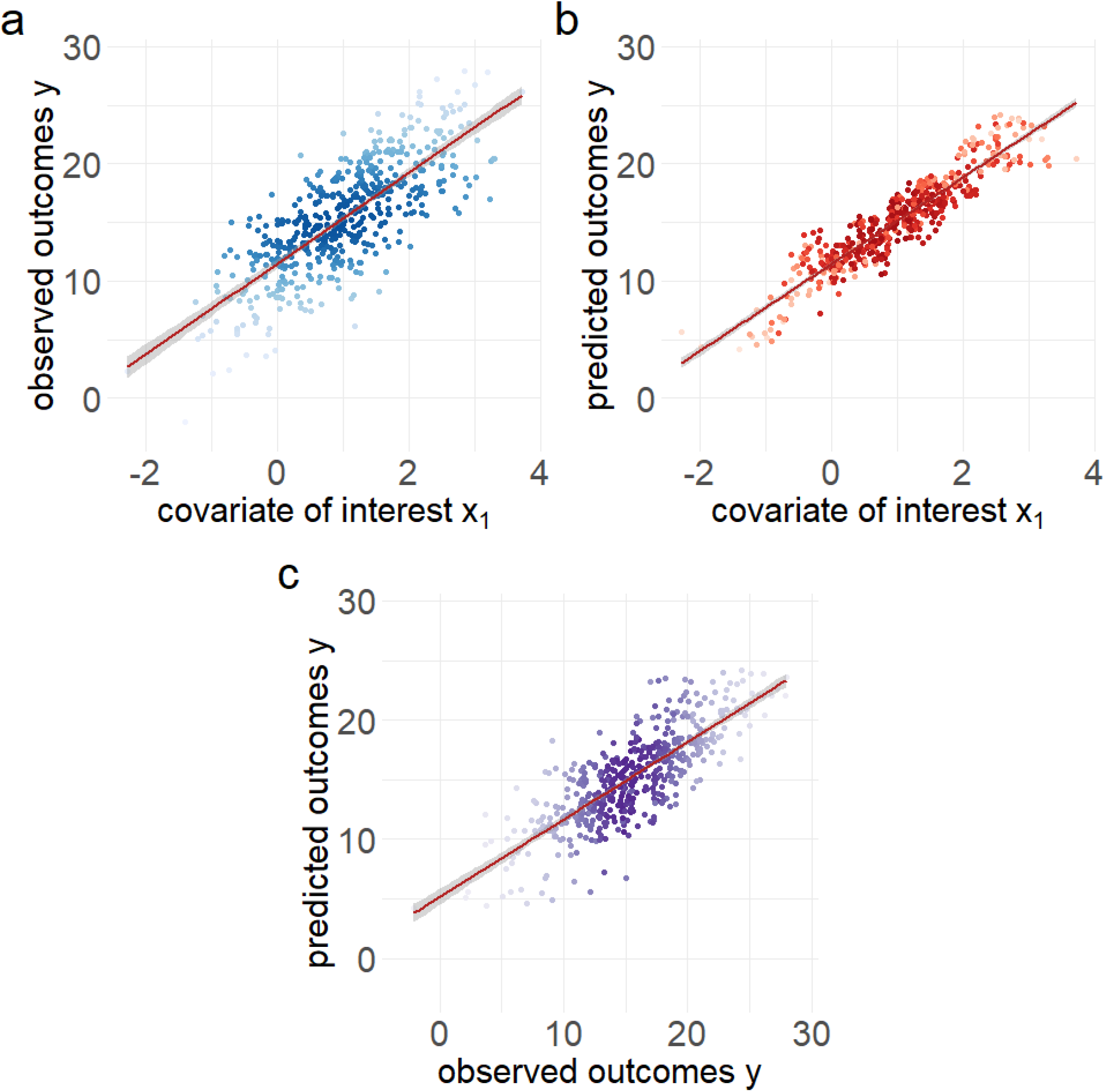
Simulated example. Data were simulated from the ground truth model as a linear model. (a) Observed outcomes versus the covariate of interest. The x-axis shows the covariate of interest *x*_1_ and the y-axis shows the observed outcomes of *y*. (b) Predicted outcomes versus the covariate of interest. The x-axis shows the covariate of interest *x*_1_ and the y-axis shows the predicted outcomes of *y_p_*. (c) Observed outcomes versus predicted outcomes. The x-axis shows the observed outcomes of *y* and the y-axis shows the predicted outcomes of *y_p_*.

This simulation is designed to highlight the issues that arise with post-prediction inference in a setting where both *y_i_* and *y_pi_* are available. In actual data analysis with predicted outcomes, we would not observe the true *y_i_* in the validation set and all inference would be performed with *y_pi_*.

In the first panel Figure 2(a) we illustrate the true relationship between the simulated *y* and *x*_1_ (blue color). In the second panel Figure 2(b) we show the predicted values *y_p_* versus *x*_1_ (red color). In this figure, the relationship has changed, with different slope and variance. In the third panel Figure 2(c) we show the relationship between the observed and predicted outcomes. In this simulated example, we know that the estimated coefficient for the relationship between the observed outcome *y* and *x* is 3.87 with a standard error of 0.14. However, when we fit model using the predicted outcome *y_p_* we get an estimate of 3.7 with a standard error of 0.068. This simple simulated example illustrates that inferences drawn with predicted outcomes may have (a) biased estimates, (b) too small standard errors, and hence (c) p-values and inference that are anti-conservatively biased.

To adjust for error in predictions, one option would be to derive bias and standard error corrections for a specific machine learning method. This approach would leverage knowledge about how a specific prediction tool works. To compute the bias and standard errors analytically we both (a) need to know what machine learning model was used and (b) need to be able to theoretically characterize the properties of that machine learning model’s predictions. This approach, would restrict an analyst to only machine learning approaches whose inferential operating characteristics have been derived. Figure 2(c) suggests an alternative approach. In this case, the relationship between the observed and predicted outcome can easily be modeled using linear regression. We will show that this observation holds for a variety of machine learning techniques.

The key idea of our approach is that we use the relationship between the predicted and observed data in the testing set, to estimate the bias and variance introduced by using predicted outcome as the dependent variable in the downstream inferential regression model in the validation set. This approach does not require idiosyncratic information about each machine learning approach and, instead, assumes that a relatively simple model captures the relationship between the predicted and observed outcomes.

## 3 Method

### 3.1 Overview of our approach

Our goal is to develop a method for correcting inference for parameters in an inferential regression model where predicted outcomes are treated as observed outcomes.

To do this we make the following assumptions about the structure of the data and model. We assume that the data are generated from an unknown data generating model of the form:

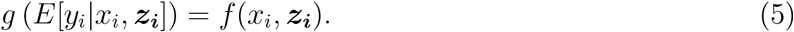

Here *x_i_* denotes the covariate of interest and *z_i_* denotes other covariates. This model represents the “true state of nature” but is not directly observed in any practical problem.

We also assume that in a new data set it may be too expensive, too time-consuming, or too difficult to collect outcome variable *y_i_* for all samples. We, therefore, will attempt to predict this outcome with an arbitrary machine learning algorithm 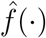 so that 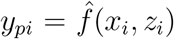 is the predicted outcome based on the observed covariates.

However, the primary goal of our analysis is not to simply predict outcomes but to perform inference on the relationship between the outcomes and covariates. This may be either the set of covariates used in training the model or new covariates collected in subsequent data sets.

In practice, the true data generating process is rarely know. Common statistical practice is to fit linear or generalized linear models to relate outcomes to covariates for inference. Letting *X* to denote the covariates of interest in matrix notation, then a typical regression model may be of the form:

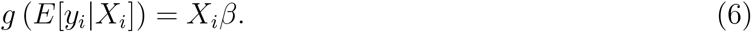

When the outcome is observed, we can directly compute the estimate of *β*. However, here we consider the case where it will not be possible to observe the outcome in future data sets due to cost or inconvenience, so the predicted outcome *y_pi_* will be used in Equation (6).

The most direct approach to performing post prediction inference is to use predicted outcomes and ignore the fact that they are predicted. However, this approach can lead to bias in the estimates, small standard errors and anti-conservatively test statistics and false positives for estimated coefficients as we saw in the simple example in Section 2.1. We will demonstrate that this approach produces consistently inaccurate inference in the simulation and real application settings. Despite these potential biases, this approach to direct use of predicted outcomes in inferential models is popular in genomics [9], genetic [6], public health [18], and EHR phenotyping [1] among other applications.

Another strategy would be to try to directly derive the properties of the coefficients and standard errors in the subsequent inference model using the definition of the machine learning algorithm *f* (·). When a prediction is based on a sufficiently simple machine learning algorithm, this may be possible to do directly. However, machine learning models now commonly include complicated algorithmic approaches involving thousands or millions of parameters, including k-nearest neighbors [32], SVM [33], random forests [30, 31] and deep neural networks [34].

We instead focus on modeling the relationship between the observed and predicted outcomes. Our key insight is that even when we use a complicated machine learning tool to predict outcomes, a relatively simple model can describe the relationship between the observed and predicted outcomes (Figure 3). We then use this estimated relationship to compute bias and standard error corrections for the subsequent inferential analyses using predicted values as the outcome variable.

**Figure 3:**
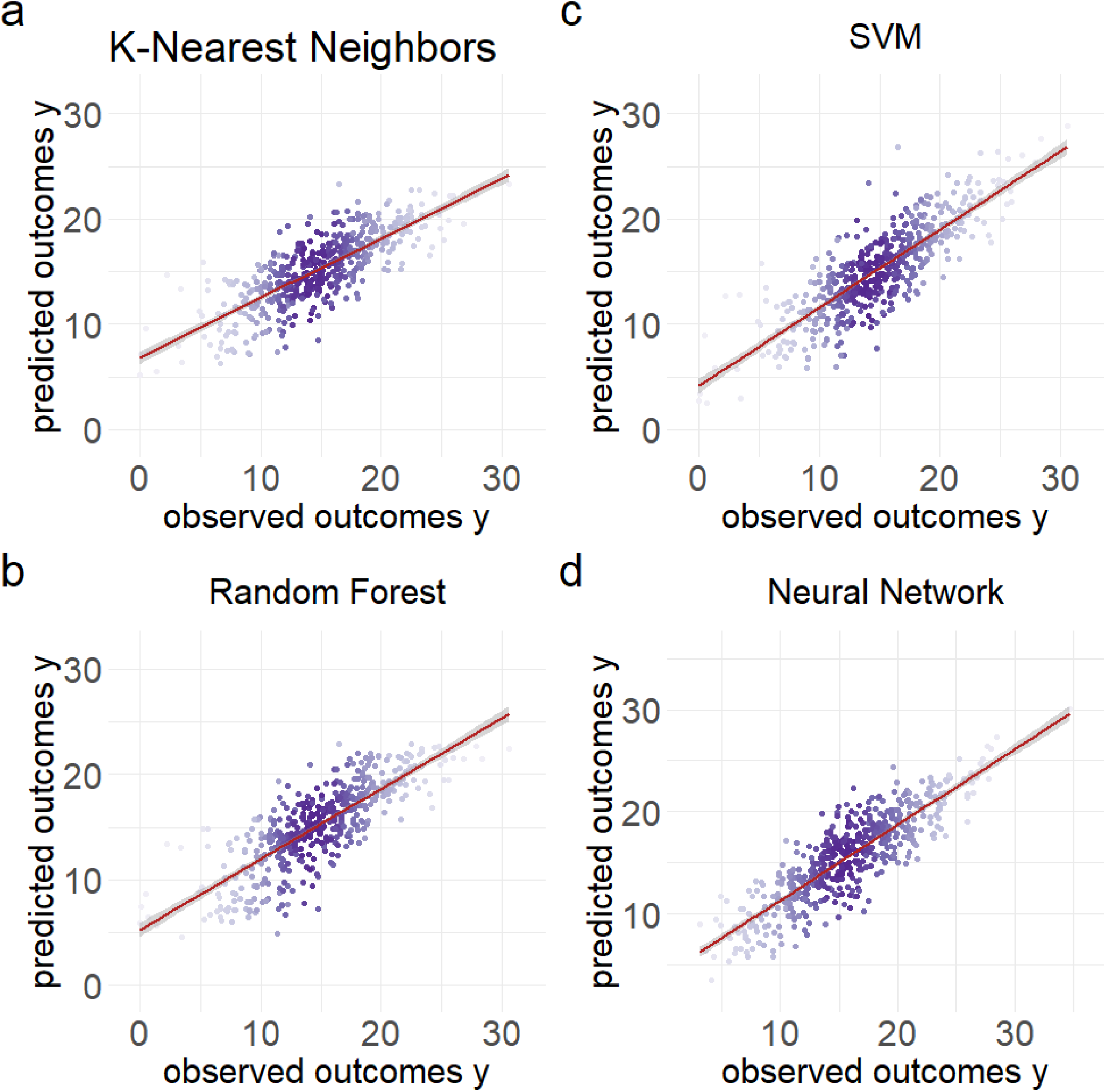
Relationship between the observed and predicted outcomes using different machine learning models. Data were simulated from the ground truth model as a linear model with normally distributed noise. On the x-axis is the observed outcome of *y* and on the y-axis are the predicted outcomes *y_p_*. We show that regardless of the prediction method (a) k-nearest neighbors, Random Forests, (c) SVM, or (d) Neural Network, the observed and predicted outcomes follow a distribution that can be accurately approximated with a regression model.

Based on the observation in Figure 3, we relate the observed to the predicted data through a flexible model *k*(·):

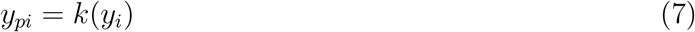

For continuous outcomes, we can estimate this relationship as a linear regression model. For categorical outcomes, we can use a logistic regression model or a simple machine learning model to estimate the relationship between the observed and predicted outcomes. To fit this relationship model we take advantage of the standard structure of machine learning model development. In these problems, the observed data is split into training, testing, and validation sets, and we assume that the three sets have the same data generating system. As illustrated in figure 1, we can build the prediction model in the training set and then compute an unbiased estimate of the relationship model in the testing set. Using this relationship model we derive a correction for the estimates, standard errors, and test statistics for our inference model. Then in the validation set, we can evaluate the quality of our correction in an independent sample.

In the following two sections, we derive bootstrap-based and analytical methods to correct inference for parameters in an inferential model on future data sets where predicted outcomes are treated as observed outcomes. For both methods, we generalize the approach to split the data into training, testing, and validation sets, and we assume that the three sets follow the same data generating procedures. With either method, we assume that the covariate of interest can be one of the covariates observed in the training and testing set, or a new covariate measured only in the validation or future data set.

In Section 3.2, we develop a flexible bootstrap procedure for post-prediction inference correction. The bootstrap based approach allows for flexibility in both the relationship model and the subsequent inferential models. This approach is applicable provided that the relationship can be modeled through any sufficiently simple relationship that allows bootstrap sampling. In Section 3.3, we derive an analytic correction that can be applied subject to additional assumptions. For the analytical derivation, we assume (1) the outcome is continuous on the training, testing, and validation set, (2) relationship between the observed and predicted outcomes can be modeled using a normal linear regression model and (3) that the inferential goal is a linear regression model on the validation set. Under these assumptions, the analytic correction holds regardless of the choice of machine learning algorithm *f* (·) used to make the predictions.

### 3.2 Bootstrap correction

In this section, we propose a bootstrap approach for correcting the bias and variance in the down-stream inferential analyses. This approach can be applied for continuous, non-normal data, categorical data, or count data. For our approach we make the following assumptions: (1) we have a training set to build the prediction model, a testing set to estimate parameters of the relationship model, and a validation set to fit a generalized regression model as the subsequent inferential model, and all three sets must follow the same data generating system, (2) the covariate of interest in the subsequent inferential analyses can be either a variable already seen on the training and testing sets, or a new variable collected only in a new sample on the validation set, (3) the relationship between the observed and predicted outcome can be modeled through a flexible but specific simple model in the form of *y_i_* = *k*(*y_pi_*) that is easy to sample from, and (4) the relationship model will hold in future samples.

The first step of our bootstrap procedure is to randomly split the data into a training, testing, and validation set as illustrated in figure 1. The algorithm then proceeds as follows:

**<u>Bootstrap Procedure:</u>**

1. Use the observed outcomes and covariates in the training set (*y*_(*tr*)_*, x*_(*tr*)_) to estimate a prediction model 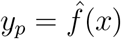.
2. Use the observed outcomes and predicted outcomes in the testing set (*y*_(*te*)_*, y_p_*_(*te*)_) to estimate the relationship model *y* = *k*(*y_p_*), where *k*(·) can be any flexible function.
3. Use the predicted outcomes and observed covariates in the validation set (*y_p_*_(*val*)_*, x*_(*val*)_) to bootstrap as follows. **for** Bootstrap iteration b = 1 to B **do**
  i. For *i* = 1, 2, …, *n*, sample predicted values and the matching covariates 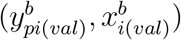 with replacement
  ii. Simulate values from the relationship model 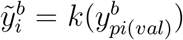 using the function *k*(·) estimated from the testing set in Step 2
  iii. Fit the inference model 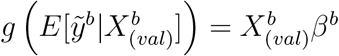 using the simulated outcomes 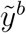 which build in the prediction error from the relationship model and the matching model matrix based on the sampled covariates in matrix notation *X^b^*
  iv. Extract the coefficient estimator 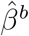 from the fitted inference model in (iii)
  v. Extract the standard error of the estimator 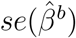 from the fitted inference model in (iii) **end for**
4. Estimate the inference model coefficient using a median function on the estimators 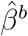 collected in Step 3(iv): 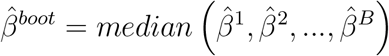.
5. Estimate the the inference model standard error:
  - For the parametric method, use a median function on the the standard errors 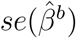 collected in Step 3(v): 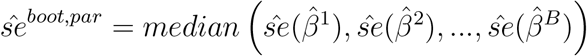
  - For the non-parametric method, use the standard error of the estimators 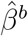 collected in Step 3(iv): 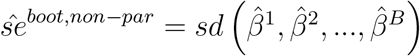

The bootstrap approach builds in two types of errors: the error due to random sampling and the prediction error. The prediction error is introduced by sampling from the relationship model in the ***for*** loop Step 3(ii). We again make the simplifying assumption that *y* and *y_p_* can be related through a model that is easy to fit. We can focus here on the class of generalized linear models, but in the ***Bootstrap Procedure*** Step 2, the relationship function *k*(·) could be more general, even flexible as a machine learning algorithm, provided it can be easily estimated and sampled. The advantage of the relationship model is that we do not need to assume the type or complexity of the function *f* (·) used to make the predictions. It can be arbitrarily complicated so long as the estimated relationship between the observed and predicted values can be sampled.

### 3.3 Analytical derivation of a correction

In this section, we propose an analytical method to correct inferences for the parameters in the linear downstream inference model. We assume that the data have been divided into training (tr), testing (te), and validation sets (val) and that the data generating distribution is the same across the three sets: 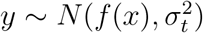 where *f* (·) is an arbitrary and unknown function of the covariates. In the training set, we use the observed outcomes and covariates (*y*_(*tr*)_*, x*_(*tr*)_) to estimate a prediction model 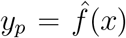. In the testing set, we use the predicted and observed outcomes (*y*_(*te*)_*, y_p_*_(*te*)_) to estimate a linear relationship model. In the validation set, we would fit a linear inference model using predicted outcomes and covariates in matrix notation (*y_p_*_(*val*)_*, X*_(*val*)_). Our goal is to infer the relationship between the outcome *y* and some subset of the covariates in the validation set or a future data set where collection of the outcome is either prohibitively expensive or complicated.

The analytical derivation approach computes the corrected parameters in the inference model more efficiently than the bootstrap approach, but with more restrictions in the assumptions to calculate a closed for solution to the parameters in the downstream inferential model: (1) we concentrate on a setting where the outcome can only be continuous and approximately normally distributed, (2) the relationship model estimated on the testing set is also approximately normally distributed, (3) the subsequent inferential model must be a linear model that we can correct inference from, and (4) the covariate of interest in the inference model can be either a variable already seen on the training and testing sets, or a new variable collected only in a new sample on the validation set.

In the validation set ideally we would fit the model

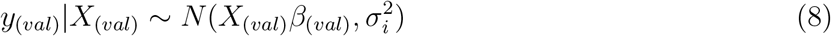

However, the outcome is not observed in the validation set. We therefore instead will fit the model:

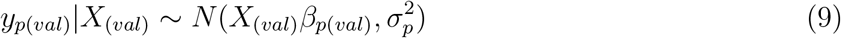

where we are no longer estimating the same quantity due to the change in dependent variable. This uncorrected strategy to post-prediction inference is commonly used in real practices [1, 5, 6, 7, 8, 9]. Our goal here is to develop a correction to recover the inference about *β*_(*val*)_ we would have made had the observed outcomes been available.

We can use information about the relationship between the observed outcome *y* and predicted outcome *y_p_* to correct inference in data sets where *y* is not observed and we substitute *y_p_*. We make the assumption that the observed outcomes and predicted outcomes follow a relationship model:

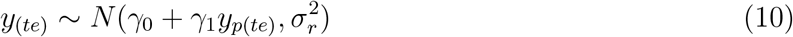

The key observation we have made is that a simplified model often holds, even when the machine learning function used to make the predictions 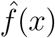 is quite complicated (see figure 3).

Our goal is not to model the full distribution of (*y*_(*val*)_, *y_p_*_(*val*)_, *x*_(*val*)_), but instead to infer the relationship between the outcome *y*_(*val*)_ and a set of covariates *x*_(*val*)_. If we had observed *y*_(*val*)_ and fit the inference model as shown in Equation (8), we have the OLS estimator 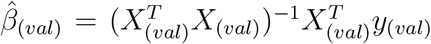. However, *y*_(*val*)_ is unobserved and thus 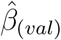 cannot be calculated directly. So, we first want to estimate *y*_(*val*)_ using the conditional expectation *E*[*y*_(*val*)_|*X*_(*val*)_]. This expectation can be written as:

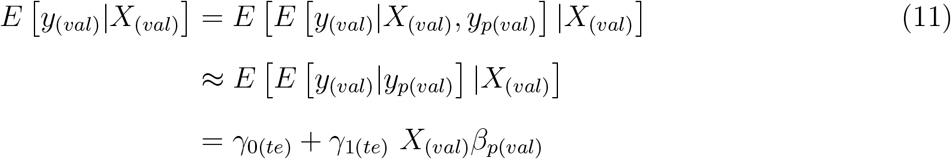

Here *β_p_*_(*val*)_ represents the parameter in the linear regression inference model where predicted outcome is used as the dependent variable. The approximation in Equation (11) is based on using the relationship between the predicted outcome and observed outcome *E*[*y*_(*val*)_|*y_p_*_(*val*)_] as an approximation to the conditional expectation *E*[*y*_(*val*)_|*X*_(*val*)_*, y_p_*_(*val*)_] (see supplement Section 1.1.1 for full analytical derivation).

This approximation can be made exact in the extreme scenario where the predicted outcome exactly captures the relationship between the outcome and the covariates *y_p_* = *f* (*x*). In this case, the real outcome can be written as *y* = *y_p_*+*ϵ*, and we have exactly *E*[*y*_(*val*)_|*X*_(*val*)_*, y_p_*_(*val*)_] = *E*[*y*_(*val*)_|*y_p_*_(*val*)_] (see supplement Section 1.1.4 for a full analytical derivation). Thus, we can approximate the unobserved outcome *y*_(*val*)_ as

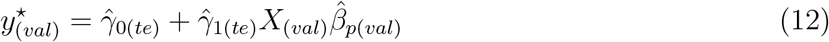

and we then approximate the estimator 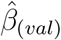 as

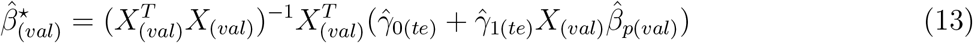

Through this approximation, we further show that 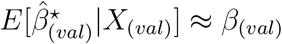 (see supplement Section 1.1.2 for full analytical derivation).

To make inferences, we also need to estimate the standard error of the estimator 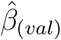. The challenge is that the standard error cannot be simply calculated by fitting the regression model in Equation (8) because *y*_(*val*)_ is unobserved. Instead, we first estimate the conditional variance *V ar*[*y*_(*val*)_ | *X*_(*val*)_] using the variance that comes from both the relationship model in Equation (10) and the inference model in Equation (9) with predicted outcomes. This is a similar approach to the expectation derivation above where we assume that the observed outcome is unknown. Using the law of total conditional variance:

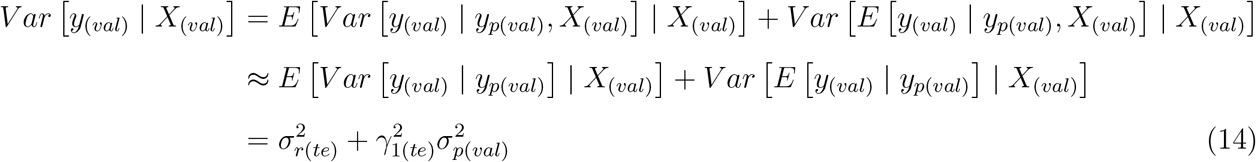

where in the second step of Equation (14), we again have made the approximation of using the relationship between *y*_(*val*)_ and *y_p_*_(*val*)_ to model the conditional variance *V ar*[*y*_(*val*)_|*y_p_*_(*val*)_*, X*_(*val*)_]. We show that under the extreme case where the predicted outcome exactly captures the relationship between the outcome and the covariates, we have exactly *V ar*[*y*_(*val*)_|*y_p_*_(*val*)_*, X*_(*val*)_] = *V ar*[*y*_(*val*)_|*y_p_*_(*val*)_] (see supplement Section 1.1.4 for full analytical derivation). Then we estimate the standard error of the estimator 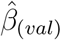 (see supplement Section 1.2 for full analytical derivation):

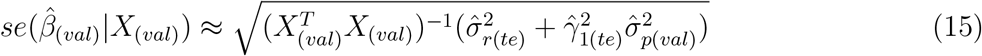

Therefore, with the estimated corrected coefficient 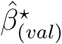 and the estimated standard error 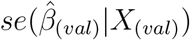, we now are able to estimate a test statistic to recover the inference we would have made in Equation (8) when the observed outcomes had been available (see supplement Section 1.2 for details in hypothesis test and defined decision rule). The test statistic is approximated as:

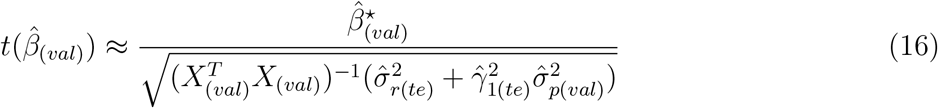

### 3.4 Simulated data

We simulate the independent covariate *x*, the error term *e_u_*, and then simulate observed outcome *y* using the “true state of nature” model in Equation 5. The “true state of nature” is not directly observed in practical problems but can be specified in simulated problems. We consider both the case of a continuous outcome in Section 3.4.1 and a binary outcome in Section 3.4.2 that demonstrate uncorrected post-prediction inference leads to bias in the estimates, small standard errors and anti-conservative test statistics.

We also include simulations that demonstrate anti-conservative bias in p-values from uncorrected post-prediction inference in Supplemental Section 2. The key insight of our postpi methods relies on the fitness of the relationship between the observed and predicted outcomes (*y* and *y_p_*) estimated in the testing set. In many cases, this relationship can be well described as a simple model but this may not always hold true. For instance, when the predicted values are obtained from weak learners, the correlation between the observed and predicted outcomes may not be sufficiently strong to allow corrected inference. As expected, we observe improved operating characteristics of our methods with increasing accuracy of the prediction model. We show that our postpi methods successfully approximate the estimates, standard errors, t-statistics, and p-values as we would have obtained using the observed *y* (Supplemental Figures 1 and Figure 2). We also show our corrections are reasonably robust to the levels of correlation between *y* and *y_p_* ranging from 0.1 - 0.2 to 0.7 - 0.8, and across all levels of correlation our postpi methods successfully correct the distribution of p-values compared to the uncorrected post-prediction inference - recovering type I error rate control (Supplemental Figure 1).

#### 3.4.1 Continuous case

For the continuous case we simulate covariates *x_ij_* and error terms 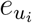 from normal distributions, and simulate the observed outcome *y_i_* using a linear function *h*(·) as the “true state of nature” model for *i* = 1, …, *n*, *j* = 1, …, *p*.

In each simulation cycle, we set the total sample size *n* = 900 and the dimension of covariate matrix *p* = 4. To mimic a complicated data generating distribution and make predictions sufficiently variable for illustration purposes, we generate data including both linear and smoothed terms. For the smoothed terms, we use Tukey’s running median smoothing with a default smoothing parameters“3RS3R” [35]. The error terms are also simulated from a normal distribution with independent variance. The model specification is:

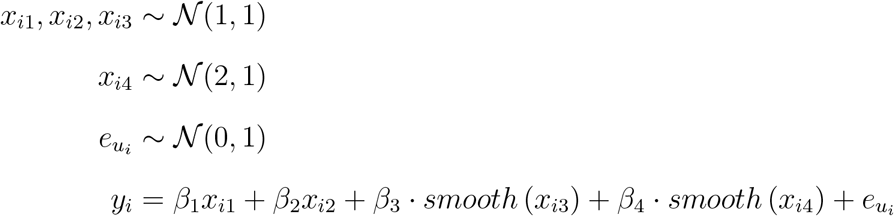

We create a training, testing and validation set by randomly sampling the observed data into three equal size groups each with sample size 300. To mimic a more realistic setting, we assume that we are only interested in associating the outcome (*y_i_*) and one covariate (in this case *x_i_*_1_), and we will use a linear inference model to quantify this relationship.

For our simulation we fit a generalized additive model (GAM) [36] to the data in the training set. To estimate the prediction function *f* (·), we use all of the covariates *x_i_*_1_*, x_i_*_2_*, x_i_*_3_*, x_i_*_4_ as features to predict the observed outcomes *y_i_*. This prediction is meant to simulate the case where we are trying to maximize predictive accuracy, not to perform statistical inference.

In the testing set we apply the trained prediction model to get predicted outcomes *y_pi_*. We estimate the relationship between the observed and predicted outcome (*y_i_* and *y_pi_*) as a simple linear regression model: 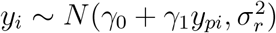. We then use standard maximum likelihood estimation to approximate the parameter estimates, 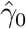, 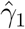 and 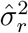.

Our evaluation of the performance of different methods is done on an independent validation set by fitting a linear regression model as the inference model. We will compare inference using the predicted outcome with no-correction, *post-prediction inference* through analytical derivation postpi and *post-prediction inference* through parametric bootstrap postpi, and non-parametric bootstrap postpi. In this simulation we also have the observed outcome *y*, so we can calculate the coefficients, estimates, and statistics that come from using the observed values in inferential models. The baseline model we are comparing to fits the regression model *E*[*y_i_*|*x_i_*_1_] = *β*_0_ + *x_i_*_1_*β*_1_ to the observed data on the validation set. We then estimate the coefficient, standard error, and t-statistic using standard maximum likelihood estimation.

To fit the three correction approaches we first perform the following steps. In the training set, we estimate the prediction function 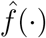. We then predict the outcome on the testing and validation sets to produce outcome predictions 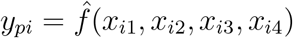.

To fit the no correction approach, we perform a regression of the form: *E*[*y_pi_*|*x_i_*_1_] = *β_p_*_0_ + *x_i_*_1_*β_p_*_1_, treating the predicted outcome as if it was observed and calculate the coefficient, standard error, and t-statistic using maximum likelihood, ignoring the fact that the outcome is predicted.

To fit the postpi analytical derivation approach, we estimate the coefficient by estimating the bias between the coefficients we get using the observed and predicted data In the testing set. We estimate the variance of this term using both the inference model and the relationship model on the testing set. We then apply these corrections to estimate the postpi analytical derivation coefficient estimator, standard error, and test statistics.

To fit the parametric and non-parametric bootstrap postpi approaches, we follow the ***Bootstrap Procedure*** Step 1-5 in Section 3.2. In Step 1 we fit the GAM [36] prediction model In the training set. In Step 2 we estimate the relationship model *y_i_* = *k*(*y_pi_*) where *k*(·) is again a GAM[36] model in the testing set. In Step 3 we first set the bootstrap size *B* = 100 to start the ***for*** loop, and then repeat Step 3(i)-(v) on the validation set. Specifically, in Step 3(ii) we estimate the relationship model *k*(·) as a linear function and simulate values from the distribution: 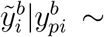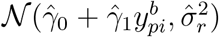. Both the mean and standard deviation of the sampling distribution come from the estimated relationship model in Step 2. We then fit a linear regression model as the inference model: 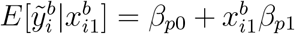 in Step 3(iii) and collect estimated bootstrap coefficient estimators and standard error in Step 3(iv)-(v). We finally estimate the coefficient in Step 4 and standard error in Step 5 for both the parametric and non-parametric bootstrap postpi methods.

Across 300 simulated cases, we fix the values of *β*_2_ = 0.5*, β*_3_ = 3*, β*_4_ = 4 and set *β*_1_ to be a range of values in [−6, −5*, …,* 5, 6] for the covariate of interest *x_i_*_1_ in the downstream inferential model. We then compute the estimates, standard errors, and t-statistics for *β*_1_ with no correction, analytical derivation postpi, parametric bootstrap postpi and non-parametric bootstrap postpi methods, and compare them to the baseline results where the outcome is observed.

We will use hextri plots to compare multiple scatter plots simultaneously [37]. These plots are designed so that the size of each bin is proportional to the number of points in the bin and they are divided into colors in proportion to the number of points from each comparison. In this simulation example, the prediction has relatively little bias, so the estimated coefficients using the predicted outcome are relatively close to the estimates using the observed outcome. In Figure 4(a) all of the colors lie close to the line of equality. However, the standard errors for the no correction approach (orange color) in Figure 4(b) is much lower than what we would have observed in the observed outcomes. This is because the prediction function attempts to capture the mean function, but not the variance in the observed outcome. We compute the root mean squared error (RMSE) [38] to show that both the postpi analytical derivation and postpi bootstrap approaches outperform the no correction approach. The standard errors are closer to the truth with an RMSE reduced from 0.088 for no correction (orange color) to 0.015 for analytical derivation postpi (green color), and also improved to 0.015 for parametric bootstrap postpi (dark blue color) and 0.019 for non-parametric bootstrap postpi (light blue color) in Figure 4(b). The improved standard errors are reflected in improved t-statistics using analytical derivation postpi and the two bootstrap postpi approaches in Figure 4(c), with RMSE reduced from 26.33 for no correction (orange color) to 2.45 for analytical derivation postpi(green color), and improved to 2.41 for parametric bootstrap postpi (dark blue color) and 2.89 for non-parametric bootstrap postpi (light blue color).

**Figure 4:**
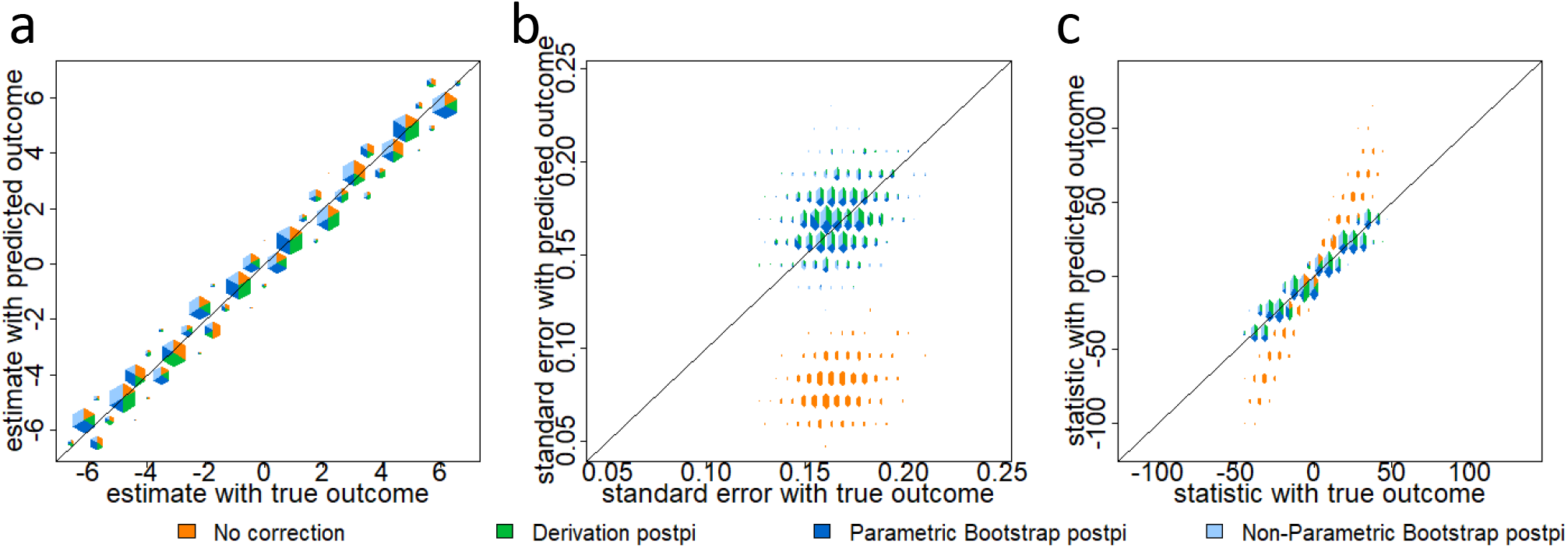
Continuous simulation. Data were simulated from the ground truth model as described in Section 3.4.1. On the x-axis are the values calculated using the observed outcome and on the y-axis are the values calculated using no correction (orange color), analytical derivation postpi (green color), parametric bootstrap postpi (dark blue color), and non-parametric bootstrap postpi (light blue color). We show (a) the estimates are similar across all four approaches since the data were simulated from a normal model, (b) the standard errors are too small for the uncorrected inference (orange color) but corrected with our approaches and (c) the t-statistics are anti-conservatively biased for uncorrected inference but corrected with our approaches.

#### 3.4.2 Binary case

For the binary case we simulate a categorical covariate *x_ic_*, continuous covariates *x_i_*_1_*, x_i_*_2_, and an error term 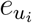, and then simulate the observed outcome *y_i_* assuming a generalized linear model *f* (·) for *i* = 1, …, *n*. In this case, we specify the “true state of nature” model *f* (·) to be a logistic regression model. To simulate observed outcomes *y_i_*, we first set up covariates through a linear combination where we smooth a subset of continuous covariates using Tukey running median smoothing [35] and include errors to increase variability in outcomes *y_i_*. We apply the inverse logit function to the linear predictor to simulate probabilities which we use to simulate Bernoulli outcomes (*y_i_* = 0 or 1) through binomial distributions. We simulate as follows:

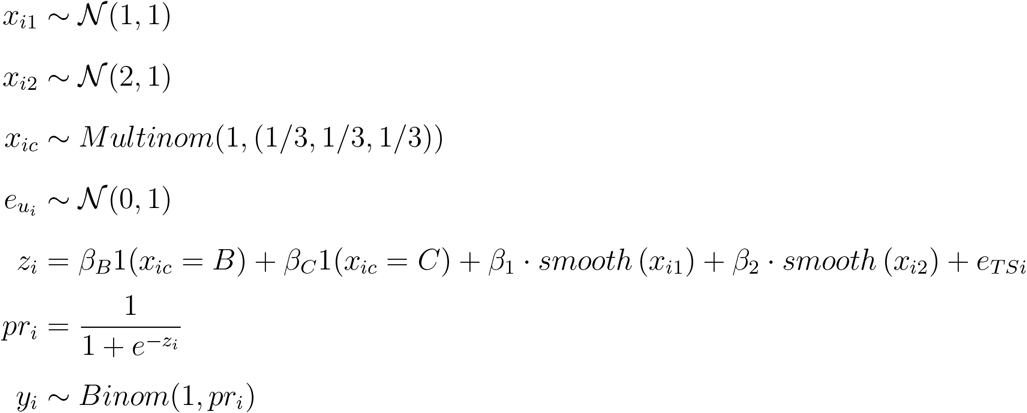

We generate 1,500 samples for each iteration and and separate the data into a training, testing and validation set of equal size n = 500. We set 1(*x_c_* = *C*) as the covariate of interest in the subsequent logistic regression inferential model. Then we use the two bootstrap methods - parametric and non-parametric bootstrap postpi to estimate the corrected coefficient estimate, standard error and test statistic through the ***Bootstrap Procedure*** Step 1-5 described in Section 3.2.

In the training set, we use a k-nearest neighbors [32] model as a machine learning tool and all independent covariates *x_ic_, x_i_*_1_*, x_i_*_2_ as features to estimate the prediction function *f* (·) on the training set. Then we apply the trained prediction model in the testing and validation sets to get the predicted outcome *y_pi_* as well as the probability *pr_i_* of the predicted outcomes (i.e. *pr_i_* = *Pr*(*y_i_* = 1)). In the testing set, we use a logistic regression to estimate the relationship between the observed outcome and the predicted probability: *g*(*E*[*y_i_* = 1|*pr_i_*]) = *γ*_0_ + *pr_i_γ*_1_, where *g*(·) is the natural log of the odds such that 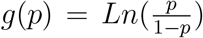. Here we form the relationship model with the predicted probability. The reason is that the outcome is dichotomous, so we have little flexibility to model the variance in the observed outcome as a function of the predicted outcome. Instead, using predicted probability provides more flexibility to model the relationship.

In the case of a categorical outcome, the analytical derivation approach no longer applies, so we apply the two bootstrap correction methods only. In the validation, set we follow the ***Bootstrap Procedure*** Step 1-5. First we set the bootstrap size *B* = 100 to start the ***for*** loop. In Step 3(ii) 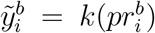, we simulate values in two steps: (1) use 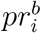 and the estimated relationship model to predict the probability of getting the “success” outcome (i.e. 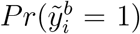), and then (2) sample 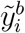 from a binomial distribution with the probability parameter as 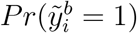 obtained from (1). In Step 3(iii) we again fit a logistic regression model as the inference model: 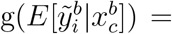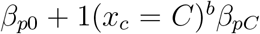. Then in Step 4-5 we estimate the parametric and non-parametric bootstrap postpi coefficient, standard error, and test statistics.

Across the 300 simulated cases, we fix the values of *β*_1_ = 1*, β*_2_ = −2*, β_B_* = 1. Here we choose 1(*x_c_* = *C*) as the covariate of interest in the downstream inferential analyses, so we set *β_C_* to be a range of values in [−2, −1.5*, …,* 4.5, 5]. Under many simulations, there is a problem of sparsity in the dichotomous covariates where inference from even observed *y_i_* would be instable. In this example, we exclude such sparse cases in the simulations which lead to extremely large standard errors and inaccurate estimates across all approaches. In both Figure 5(a) and (b), we see that the estimates and standard errors are inflated in the case of no correction (orange color). In detail, we see bias in the coefficient estimate using the no correction approach (orange color) in Figure 5(a) with RMSE 2.94 compared to the truth. This bias is corrected through the parametric bootstrap (dark blue color) and non-parametric bootstrap (light blue color) postpi methods with RMSE reduced to 0.53. The standard errors for no correction (orange color) in Figure 5(b) have RMSE 0.49 but reduced to 0.018 for parametric bootstrap postpi (dark blue color), and 0.025 for non-parametric bootstrap postpi (light blue color). In Figure 5(c), the t-statistics have RMSE 2.06 using no correction (orange color), 2.04 for parametric bootstrap postpi (dark blue color), and to 2.12 for non-parametric bootstrap postpi (light blue color). We observe a slight conservative bias in the t-statistics due to the postpi corrections - the blue points are consistently slightly below the line of equality. This conservative bias is an acceptable tradeoff in cases where the observed outcomes are not available.

**Figure 5:**
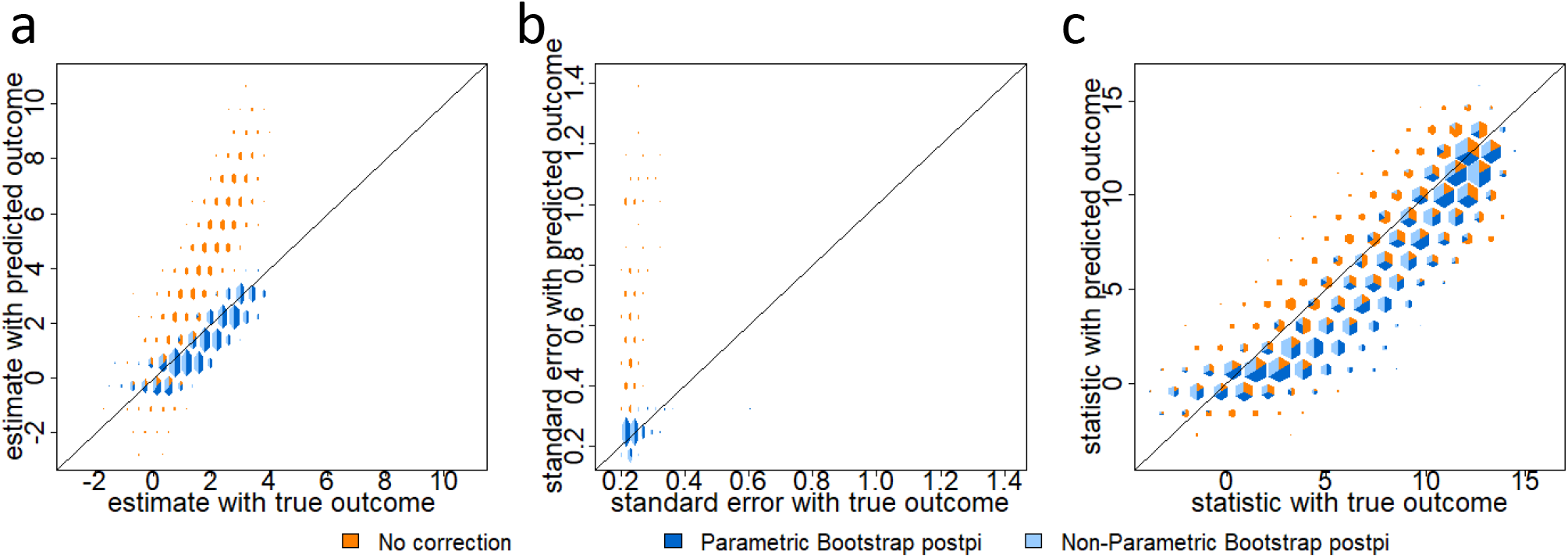
Categorical simulation. Data were simulated from the ground truth model as described in Section 3.4.2. On the x-axis are the values calculated using the observed outcome and on the y-axis are the values calculated using no correction (orange color), parametric bootstrap postpi (dark blue color), and non-parametric bootstrap postpi (light blue color). We show (a) the uncorrected estimates are anti-conservatively biased but this bias is corrected with our postpi approaches, (b) the uncorrected standard errors are also inflated and corrected by postpi and (c) the t-statistics show a slight conservative bias compared to the no-correction case.

### 3.5 Applications

To demonstrate the wide applicability of our methodology for performing *post-prediction inference*, we present two examples from very different fields: genomics and verbal autopsy analysis. These applications share very little in common scientifically, but represent two high profile examples where inference is typically performed with uncorrected predictions as the outcome (dependent) variable.

First, consider the “Recount2“ Project (https://jhubiostatistics.shinyapps.io/recount) [39] which consists of RNA sequencing (RNA-seq) gene expression data for over 70,000 human samples aligned using a common pipeline processed in Rail-RNA [40]. While “Recount2“ human samples have available gene expression information, not all samples contain observed phenotype information since the majority of the samples are pulled directly from public data on the sequence read archive [41]. However, we previously showed that many of these missing phenotype data can be predicted from the genomic measurements [2]. Our goal is to perform inference using these predicted phenotypes.

Second, we describe the distribution of (predicted) causes of death. In regions of the world where routine monitoring of births and deaths is not possible, one approach to estimating the distribution of deaths by cause is the verbal autopsy (VA) survey. These surveys take place with a caregiver or relative of the decedent and ask about the circumstances surrounding the person’s death, and typically take place when deaths happen outside of hospitals or routine medical care. Either expert guidance about the relationship between reported symptoms prior to death and the eventual cause or small “gold standard” datasets are used to train algorithms that predict causes of death based on reported symptoms. Algorithm development to predict causes of death is an active area of research and is challenging since data typically contain a mixture of binary, continuous, and categorical symptoms and many causes of death have similar presentations. After assigning a predicted cause of death, a common task is to describe patterns in the cause of death distribution. A scientist may be interested, for example, in how the distribution of deaths varies by region or by sex.

#### 3.5.1 Predicting tissue types

We consider a motivating problem from the “Recount2“ Project [39] (https://jhubiostatistics.shinyapps.io/recount/). In this example, the phenotype we care about is the tissue type where the RNA is sampled from. Understanding gene expression levels across tissues and cell types have many applications in basic molecular biology. Many research topics concentrate on finding which genes are expressed in which tissues, aiming to expand our fundamental understanding of the origins of complex traits and diseases [42, 43, 44, 45, 46]. The Genotype-Tissue Expression (GTEx) project [47], for example, studies how gene expression levels are varied across individuals and diverse tissues of the human body for a wide variety of primary tissues and cell types [42, 47]. Therefore, to better understand the cellular process in human biology, it is important to study the variations in gene expression levels across tissue types.

Even though tissue types are available in GTEx [47], they are not available for most samples in the “Recount2“. In a previous paper [2], we developed a method to predict for those missing phenotypes using gene expression data. In this example, we collected a subset of samples that we have observed tissue types as breast or adipose tissues. We also had predicted values for the above samples calculated in a previous training set [2] using the 2281 expressed regions [10] as predictors. Our goal in this example is to understand which of these regions are most associated with breast tissue in new samples (i.e. samples without observed tissue types) so that we can understand which measured genes are most impacted by the biological differences between breast and adipose tissues. Although here the phenotype we care about is the tissue types, especially breast and adipose tissues, our method can be broadly applied to any predictions to all phenotypes.

To test our method, we collected 288 samples from the “Recount2“ with both observed and predicted tissue types. Among the observed tissue types, 204 samples are observed as adipose tissues and 84 samples are observed as breast tissues. The predicted values obtained from a previously trained data set [2] include the predicted tissue type (i.e. adipose tissue or breast tissue) and the probability for assigning a predicted tissue type. In this example, we compare no correction and postpi bootstrap approaches only since the outcomes we care about - tissue types are categorical.

The inference model we are interested in is: 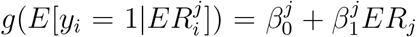. Here *g*(·) is the logit link function for *j* = 1*, . . .,* 2281 (expressed regions) and *i* = 1*, . . ., n*, *n* is the total number of samples in the “Recount2“. In the model, *y_i_* = 1 or *y_i_* = 0 represents whether breast tissue is observed or adipose tissue is observed at the *i*th sample, and 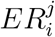 is the gene expression level for the *j*th region on the *i*th sample.

For this dataset (288 samples), we have binary tissue type outcomes. Since the predicted outcomes were obtained in a previously trained set [2], we only need to separate our data into a testing and validation set, each with a sample size *n* = 144. On the testing set, we fit a k-nearest neighbors [32] model to estimate the relationship between the observed tissue type and the probability of assigning the predicted value. On the validation set, we follow the ***Bootstrap Procedure*** in Section 3.2. Particularly in Step 3(ii), we simulate values from a distribution 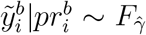. Similar to we did with the simulated data in Section 3.4, in this example, we set *F_γ_* to be a binomial distribution with the probability parameter (i.e. probability of assigning the outcome as breast cancer) estimated from the relationship model. In this way, we utilize the estimated relationship to account for necessary variations in simulated outcomes.

Among the 2281 expressed regions [10] used to make tissue type predictions [2], we care about the regions that have expression values across a relatively large amount of samples on the validation set. It is a well-known phenomenon that many RNA-seq measurements may be zero if the number of collected reads is low. To avoid highly variable model fits due to zero variance covariates, we only fit logistic regressions inference models to each filtered expressed region with expressed values over at least 20% samples. Under this filtering procedure, we include 101 expressed regions as regression variables, and fit the inference model described above to each of the region on the validation set. We then get 101 estimates, standard errors and t-statistics. We compare them to the no correction approach as we did with the simulated data.

By comparing RMSE, we observed that the estimates, standard errors and test statistics are improved from no correction to parametric and non-parametric bootstrap postpi methods. In Figure 6(a), no correction (orange color) estimates have RMSE 0.36 compared to the truth and reduces to 0.08 with parametric bootstrap postpi (dark blue color), and non-parametric bootstrap postpi (light blue color). The standard errors in Figure 6(b) have RMSE 0.08 for no correction (orange color), but corrected to 0.01 for parametric bootstrap postpi (dark blue color) and 0.03 for non-parametric bootstrap postpi (light blue color). The resulting t-statistics are improved from RMSE 0.91 for no correction (orange color) to 0.63 for parametric bootstrap postpi (dark blue color), and 0.93 for non-parametric bootstrap postpi (light blue color).

**Figure 6:**
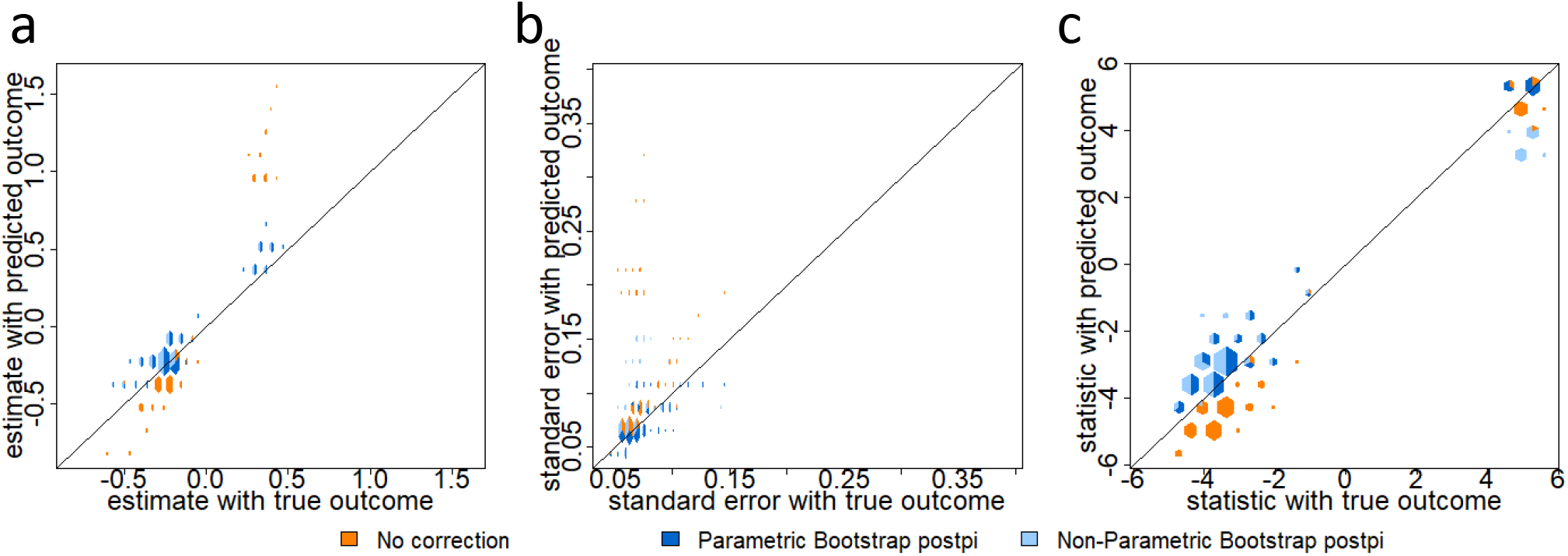
Breast versus adipose tissue prediction. Data were collected from the “Recount2” as described in Section 3.5.1. On the x-axis are the values calculated using the observed outcome and on the y-axis are the values calculated using no correction (orange color), parametric bootstrap postpi (dark blue color), and non-parametric bootstrap postpi (light blue color). We show (a) the estimates, (b) the standard errors and (c) the t-statistics. The two bootstrap postpi approaches clearly improve the estimates and standard errors compared to no correction.

We also applied our approach to correct inference for models using predicted RNA-quality as an example of how to do post prediction inference for continuous outcomes (see supplement Section 3.1).

#### 3.5.2 Describing cause of death distributions

We now move to our second example where the outcome of interest is the (predicted) cause of death and inputs are symptoms or circumstances reported by a caregiver or relative. Symptoms might include, for example, whether a person had a fever before they died, how long a cough lasted (if one was reported), or the number of times they visited a medical professional. We use data from the Population Health Metrics Research Consortium (PHMRC), which consists of about 7,800 “gold standard” deaths from six regions around the world. These data are rare because they contain both a physical autopsy (including pathology and diagnostic testing) and a verbal autopsy survey. Typically, only a small fraction of deaths will have an assigned cause (e.g. by a clinician reading the verbal autopsy survey) and these few labeled deaths will be used as inputs to train a model for the remaining deaths.

We split the data into a training and testing set, with 75% of the data used for training. The PHMRC data classify cause of death at several levels of granularity. For our experiments, we combined causes into twelve broad causes of death (Cancers, Diabetes, Renal diseases, Liver diseases, Cardiovascular causes, Stroke, Pneumonia, HIV/AIDS or Tuberculosis, Maternal causes, External causes, Other communicable diseases, and Other non-communicable diseases). We predicted the cause of death using *InSilicoVA*[11] which uses a Naive Bayes classifier embedded in a Bayesian framework to incorporate uncertainty between cause classifications.

In this example, we want to understand trends in the twelve combined causes of death and we care about both continuous and categorical symptoms. Continuous symptoms include age, number of people living at this address, age of the respondent. Categorical symptoms include year of death (2007, 2008, 2009, 2010), sex (male or female), death certificate issued (yes or no), used tobacco (yes or no), used alcohol (yes or no), education of deceased (College or Higher, High School, Primary School, No Schooling), separate room for cooking (yes or no), sex of respondent (male or female), education of respondent (College or Higher, High School, Primary School, No Schooling), region (AP, Bohol, Dar, Mexico, UP). The inference model we are interested in is: 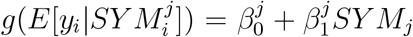. Here *g*(·) is the logit link function for *j* = 1, . . ., 13 (symptoms) and *i* = 1, . . ., *n*, *n* is the total number of samples in the dataset. In this model, *y_i_* represents one of the twelve combined causes at the *i*th sample and 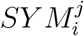 is the *j*th symptom of interest on the *i*th sample.

For this dataset, we use categorical outcomes as the causes of death for the 1960 samples and assume the outcomes are unobserved, as they typically would be in practice, for the remaining cases. Since the predicted values were obtained in a previously trained set using *InSilicoVA*[11], we only separate our data into a testing and validation set, each with a sample size *n* = 980. On the testing set, we fit a k-nearest neighbors model [32] to estimate the relationship between the observed cause of death and the probability of assigning the cause. On the validation set, we follow the ***Bootstrap Procedure*** described in Section 3.2. Particularly in Step 3(ii), we simulate values from a distribution 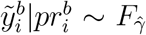. In this example, we set *F_γ_* to be a multinomial distribution with the probability parameters (i.e. probability of assigning each of the twelve broad causes of death) estimated from the relationship model as we did in the simulated data.

Among all the symptoms used to make causes of death prediction [11], we care about a subset of symptoms that also have balanced classes across the twelve broad causes of death to avoid highly variable model fits due to zero variance covariates. We then filter 13 symptoms we are interested in as regression variables and fit a logistic regression inference model to each of the selected symptom on the validation set. Because we include categorical regression variables with multiple factor levels in the inference model and get a inference result for each factor level, we obtain more factor level results than the number of symptoms. In total, we get 22 estimates, standard errors and t-statistics on the validation set. We then compare them to the no correction approach as we did with the simulated data.

We observed that the estimates, uncorrected standard errors and t-statistics (orange color) have higher RMSE compared to parametric bootstrap postpi method (dark blue color). In Figure 7(a) the RMSE of no correction (orange color) is 0.84 compared to the truth, and reduces to 0.42 with parametric (dark blue color) and non-parametric (light blue color) bootstrap postpi methods. The standard errors in Figure 7(b) have a RMSE of 0.2 for no correction (orange color), but corrected to 0.08 for parametric bootstrap postpi (dark blue color). The non-parametric bootstrap postpi (light blue color) have a RMSE 0.9, which is larger than the no correction method. In this specific case, the non-parametric bootstrap performs poorly due to imbalance in the classes leading to mis-estimated standard errors in some samples. This is a well known issue with the bootstrap for sparse outcomes [48] and would impact any bootstrap based approach for these data. The resulting t-statistics in Figure 7(c) are improved from 1.66 for no correction (orange color) to 1.24 for parametric bootstrap postpi (dark blue color). The non-parametric bootstrap postpi (light blue color) has RMSE 1.7.

**Figure 7:**
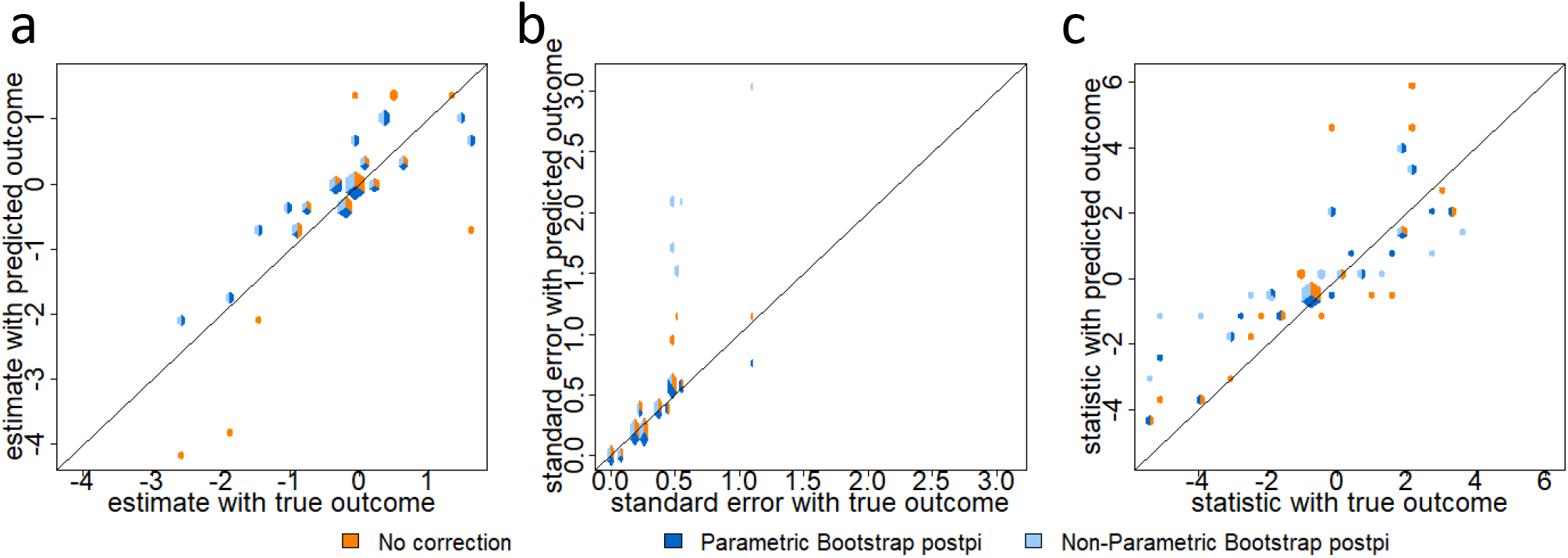
Twelve causes of death prediction. Data were collected from Population Health Metrics Research Consortium (PHMRC) described in Section 3.5.2. On the x-axis are the values calculated using the observed outcome and on the y-axis are the values calculated using no correction (orange color), parametric bootstrap postpi (dark blue color), and non-parametric bootstrap postpi (light blue color). We show (a) the estimates, (b) the standard errors and (c) the t-statistics. The parametric bootstrap postpi approach improves the RMSE of estimates, standard errors and t-statistics compared to no correction.

## 4 Discussion

As machine learning becomes more common across a range of applications, predicted outcomes will be used more often an as outcome variables in subsequent statistical analyses. As we have shown, uncorrected post-prediction inference can lead to highly variable or biased estimates of parameters of interest, standard errors that are too small, and anti-conservatively biased p-values and false positives.

We introduced methods to correct for post-prediction inference and adjust point and interval estimates when using predicted outcomes in place of observed outcomes. Our method is flexible enough to be applied to continuous and categorical outcome data, observed in fields such as medicine, public health, and sociology. Through simulated and real data, we show that our results outperform the most common current approach of ignoring the prediction step and performing inference without correction. By appropriately modeling the variability and bias due to the prediction step, the estimates, standard errors test statistics, and p-values are corrected towards the gold standard analysis we would obtain if we used the true outcomes.

Our approach relies on the key observation that the relationship between the observed and predicted values can be described as a simple model. While this observation is empirically true for the models and algorithms we considered, it may not hold universally. One limitation of our approach is that it depends on the fitness of the relationship model. For instance, when the predicted values are obtained from weak learners, the correlation between the observed and predicted outcomes is not strong, which may not be well captured by a simple model. Another limitation is that we assume the training, testing and validation sets follow the same data generating distribution. If this assumption does not hold, inference performed on the bootstrapped values on the validation set will no longer reflect the true underlying data generating process. A potential solution is that we should first conduct data normalization using methods such as SVA [49], RUV [50] and removeBatchEffect in limma [51] to correct for latent confounders in the testing or validation sets. The normalized samples can then be input into our method for subsequent inferential analyses.

Despite these limitations, we believe correction for *post-prediction inference* is crucial for obtaining accurate inference when using outcomes produced by machine learning methods. Our correction represents the first step toward a general solution to the post-prediction inference problem. To make this method usable by the community we have released the *postpi* R package: [https://github.com/leekgroup/postpi].

## Supporting information

Post-prediction inference supplementary text

## 5 Acknowledgements

Research reported in this publication was supported by the National Institute of General Medical Sciences of the National Institutes of Health under award number R01GM121459 and the National Institute of Mental Health of the National Institutes of Health under award number DP2MH122405.

## Notes

https://github.com/leekgroup/postpi

### Competing Interest Statement

The authors have declared no competing interest.

### Summary of Updates

This is a significant revision to update methodology, include new simulations, and improve flow.

