## Supplementary material for "Post-prediction Inference": Post-prediction inference supplementary text

#### 1 Analytical derivation

In the following analytical derivations we denote observed outcomes as $y$, predicted outcomes as $y_p$, covariates as $x$, and covariates in matrix notation as $X$. We assume that the data have been divided into training (tr), testing (te), and validation sets (val) and that the data generating distribution is the same across the three sets: $y \sim N(f(x), \sigma_t^2)$ where $f(\cdot)$ is an arbitrary and unknown function of the covariates. Throughout the supplement we use abbreviations for the testing, training, and validation sets when values are specific to those sets but omit them when the statements apply to all three. We also keep subscripts on $\beta$ to differentiate between sets but throughout assume $\beta = \beta_{(val)} = \beta_{(te)} = \beta_{(tr)}$. The estimated values of these coefficients, however, do differ across the three sets. We are interested in inference for the simplified model: $y \sim N(X\beta, \sigma^2)$. This represents common statistical practice where the complicated underlying functional relationship between the outcome and covariates is unknown but a regression modeling strategy is employed to perform inference. Under our problem setup the observed data consist of:

- The observed outcomes and covariates in the training set $(y_{(tr)}, x_{(tr)})$
- A prediction function estimated based on the training data $y_{p(tr)} = \hat{f}(x)$.
- The observed outcomes, predicted outcomes, and observed covariates in the test set $(y_{(te)}, y_{p(te)}, x_{(te)})$

- The predicted outcomes and observed covariates in matrix notation in the validation set and beyond $(y_{p(val)}, X_{(val)})$

Our goal is to infer the relationship between the outcome $y$ and some subset of the covariates in the validation set or a future data set where collection of the outcome is either prohibitively expensive or complicated.

In this validation set ideally we would fit the model

$$y_{(val)}|X_{(val)} \sim N(X_{(val)}\beta_{(val)}, \sigma_i^2). \tag{1}$$

However, the outcome is not observed in the validation set. We therefore instead will fit the model

$$y_{p(val)}|X_{(val)} \sim N(X_{(val)}\beta_{p(val)}, \sigma_p^2) \tag{2}$$

where $\beta_{p(val)}$ corresponds to the coefficient estimated when using the predicted value of $y$ rather than the observed. Our goal is to recover the inference we would have made had the observed outcomes been available.

As we point out in the main text in Section 3.3 we can use information about the relationship between the observed outcome $y_{(te)}$ and predicted outcome $y_{(te)p}$ in the testing set to correct inference in data sets where $y_{(val)}$ is not observed and we substitute $y_{p(val)}$. We make the assumption that the observed outcomes and predicted outcomes follow a relationship model

$$y_{(te)} \sim N(\gamma_0 + \gamma_1 y_{p(te)}, \sigma_r^2). \tag{3}$$

We can estimate the coefficients from the testing set where both $y_{(te)}$ and $y_{p(te)}$ are available and then assume that the relationship also holds in the validation set where we only have access to $y_{(val)p}$. The key insight we have made is that a simplified model often holds, even when the machine learning function used to make the predictions $\hat{f}(x)$ is quite complicated (see main text Figure 3).

#### 1.1 Conditional expectation and variance

##### 1.1.1 Approximating the conditional distribution of y

In the validation set we have assumed that only the covariates $(x_{(val)})$ and predicted outcomes $(y_{p(val)})$ are available. Critically $y_{(val)}$ is not available. As a first step, we propose an approximation of the conditional expectation of the unobserved outcome $y_{(val)}$ given covariates of interest in matrix notation $X_{(val)}$ and the predicted outcomes $y_{p(val)}$ by leveraging the assumed relationship model

$y \sim N(\gamma_0 + \gamma_1 y_p, \sigma_r^2)$ and the linear inferential model based on using the predicted outcomes $y_p \sim N(X\beta_p, \sigma_p^2)$. We approximate the conditional distribution of the unobserved $y_{(val)}$ as follows:

$$\begin{aligned}
E\left[y_{(val)}|X_{(val)}\right] &= E\left[E\left[y_{(val)}|X_{(val)}, y_{p(val)}\right]|X_{(val)}\right] && (4)\\
&\approx E\left[E\left[y_{(val)}|y_{p(val)}\right]|X_{(val)}\right] && (5)\\
&= E\left[\gamma_{0(val)} + \gamma_{1(val)} y_{p(val)}|X_{(val)}\right] && (6)\\
&= \gamma_{0(val)} + \gamma_{1(val)} E\left[y_{p(val)}|X_{(val)}\right] && (7)\\
&= \gamma_{0(val)} + \gamma_{1(val)}\ X_{(val)}\beta_{p(val)} && (8)
\end{aligned}$$

where the approximation in equation (5) is based on using the relationship between the predicted outcome and observed outcome $E[y_{(val)}|y_{p(val)}]$ as an approximation to the conditional expectation $E[y_{(val)}|X_{(val)}, y_{p(val)}]$. Since the predicted values $y_{p(val)}$ and the design matrix $X_{(val)}$ are assumed to be observed in the validation set, we can estimate $\beta_{p(val)}$ using the validation data. However, estimation of $\gamma_0, \gamma_1$ requires both the observed and predicted outcomes. By assumption, we observe both of these values in the testing set and we can therefore write:

$$\begin{aligned}
E[y_{(val)}|X_{(val)}] &\approx \gamma_{0(val)} + \gamma_{1(val)}\ X_{(val)}\beta_{p(val)} && (9)\\
&= \gamma_{0(te)} + \gamma_{1(te)}\ X_{(val)}\beta_{p(val)} && (10)
\end{aligned}$$

Where the equality in equation (10) follows from the assumption that the data generating distribution in the testing and validation sets are the same.

##### 1.1.2 Approximating the coefficients

We want to fit a linear regression model as the inferential model between $y_{(val)}$ and $X_{(val)}$. This model can be written as $y_{(val)} \sim N(X_{(val)}\beta_{(val)}, \sigma_i^2)$. In the validation set, we assume that the true outcomes $y_{(val)}$ are unobserved, our goal is to correctly estimate $\beta_{(val)}$ using the relationship model in the testing set, predicted outcome and covariates in the validation set.

If we had observed $y_{(val)}$, we have the OLS estimator $\hat{\beta}_{(val)} = (X_{(val)}^T X_{(val)})^{-1} X_{(val)}^T y_{(val)}$. Using the conditional expectation for $y_{(val)}$ we have previously computed and the relationship model from the testing set, we have $E[y_{(val)}|X_{(val)}] \approx \gamma_{0(te)} + \gamma_{1(te)} X_{(val)}\beta_{p(val)}$, and thus we can estimate the unobserved outcome $y_{(val)}$ as

$$y_{(val)}^{\star} = \hat{\gamma}_{0(te)} + \hat{\gamma}_{1(te)} X_{(val)} \hat{\beta}_{p(val)} \qquad (11)$$

Therefore, we can approximate the estimator $\hat{\beta}_{(val)}$ as

$$\hat{\beta}^{\star}_{(val)} = (X^T_{(val)}X_{(val)})^{-1}X^T_{(val)}(\hat{\gamma}_{0(te)} + \hat{\gamma}_{1(te)}X_{(val)}\hat{\beta}_{p(val)}) \tag{12}$$

The bias of this estimator can be computed as:

$$E\left[\hat{\beta}^{\star}_{(val)} - \beta_{(val)}|X_{(val)}\right] = E\left[(X^T_{(val)}X_{(val)})^{-1}X^T_{(val)}(\hat{\gamma}_{0(te)} + \hat{\gamma}_{1(te)}X_{(val)}\hat{\beta}_{p(val)})|X_{(val)}\right] - \beta_{(val)} \tag{13}$$

$$= (X^T_{(val)}X_{(val)})^{-1}X^T_{(val)}E\left[y^*_{(val)}|X_{(val)}\right] - \beta_{(val)} \tag{14}$$

where $y^*_{(val)}$ is our approximation to $y$. This expectation is a complicated function of the training, testing, and validation sets, but for good prediction functions will have expectation approximately equal to $y_{(val)}$ - which would lead to nearly unbiased estimation of $\beta_{(val)}$. We observe this behavior in the simulated examples in Section 2 of this supplement.

##### 1.1.3 Conditional variance

The analytical derivation of the conditional variance of unobserved outcome $y_{(val)}$ given covariate of interest $X_{(val)}$ in the validation set can be estimated using the variance that comes from both the relationship model $y \sim N(\gamma_0 + \gamma_1 y_p, \sigma_r^2)$ and the linear inferential model $y_p \sim N(X\beta_p, \sigma_p^2)$ using a similar approach to the analytical derivation above where we assume that the observed outcome is unknown. Using the law of total conditional variance:

$$Var\left[y_{(val)} \mid X_{(val)}\right] = E\left[Var\left[y_{(val)} \mid y_{p(val)}, X_{(val)}\right] \mid X_{(val)}\right] + Var\left[E\left[y_{(val)} \mid y_{p(val)}, X_{(val)}\right] \mid X_{(val)}\right] \tag{15}$$

$$\approx E\left[Var\left[y_{(val)} \mid y_{p(val)}\right] \mid X_{(val)}\right] + Var\left[E\left[y_{(val)} \mid y_{p(val)}\right] \mid X_{(val)}\right] \tag{16}$$

$$= E\left[\sigma^2_{r(val)} \mid X_{(val)}\right] + Var\left[\gamma_{0(val)} + \gamma_{1(val)}y_{p(val)} \mid X_{(val)}\right] \tag{17}$$

$$= \sigma^2_{r(val)} + \gamma^2_{1(val)}Var\left[y_{p(val)} \mid X_{(val)}\right] \tag{18}$$

$$= \sigma^2_{r(val)} + \gamma^2_{1(val)}\sigma^2_{p(val)} \tag{19}$$

Where the approximation in Equation (16) is the same as the conditional expectation estimated above.

Since the observed $y_{(val)}$ is not available in the testing set we can approximate the variance by borrowing estimates from the testing set for the $\gamma$ coefficients:

$$Var\left[y_{(val)} \mid X_{(val)}\right] \approx \sigma^2_{r(val)} + \gamma^2_{1(val)}\sigma^2_{p(val)} \tag{20}$$

$$= \sigma^2_{r(te)} + \gamma^2_{1(te)}\sigma^2_{p(val)} \tag{21}$$

where the equality in Equation (21) follows from the assumption that the data generating distribution in the testing and validation sets are the same.

##### 1.1.4 Extreme case

Under the extreme case where the predicted outcome exactly captures the relationship between the outcome and the covariates $y_p = f(x)$ then the real outcome can be written as $y = y_p + \epsilon = f(x) + \epsilon$. In this setting we can show that the approximation based on the relationship model in the conditional expectation analytical derivation in section 1.1.1 and the conditional variance analytical derivation in section 1.1.3 can be replaced with equality. In detail, we want to show that under the extreme case, $E(E(y|y_p, X)|X) = E(E(y|y_p)|X)$, $E(Var(y|y_p, X)|X) = E(Var(y|y_p)|X)$, and $Var(E(y|y_p, X)|X) = Var(E(y|y_p)|X)$.

Consider the simplest case where $f(X) = X\beta$, then $y = X\beta + \epsilon$, $y_p = X\beta$. We also assume $\epsilon \sim N(0, \sigma^2)$, $X \sim N(\delta, \Sigma)$. Then, we write the assumptions as follows,

$$\begin{aligned} y|X &\sim N(X\beta, \sigma^2) \\ y_p|X &\sim N(X\beta, 0) \\ X &\sim N(\delta, \Sigma) \end{aligned}$$

The marginal distribution of $y$ and $y_p$ is written as:

$$\begin{aligned} y &\sim N(\delta\beta, \beta^T\Sigma\beta + \sigma^2) \\ y_p &\sim N(\delta\beta, \beta^T\Sigma\beta) \end{aligned}$$

Therefore, $y$ and $y_p$ can be written as a bivariate normal distribution,

$$\begin{pmatrix} y \\ y_p \end{pmatrix} \sim N\left(\begin{pmatrix} \delta\beta \\ \delta\beta \end{pmatrix}, \begin{pmatrix} \beta^T\Sigma\beta + \sigma^2 & \rho(\beta^T\Sigma\beta + \sigma^2)^{1/2}(\beta^T\Sigma\beta)^{1/2} \\ \rho(\beta^T\Sigma\beta + \sigma^2)^{1/2}(\beta^T\Sigma\beta)^{1/2} & \beta^T\Sigma\beta \end{pmatrix}\right).$$

Here $\rho$ is the correlation between $y$ and $y_p$, and $\rho = \frac{(\beta^T\Sigma\beta)^{1/2}}{(\beta^T\Sigma\beta+\sigma^2)^{1/2}}$. To see this note that

$$\begin{aligned} Cov(y, y_p) &= E(yy_p) - E(y)E(y_p) \\ &= E(y_p^2 + \epsilon y_P) - (\delta\beta)^2 \\ &= E(y_p^2) - (\delta\beta)^2 \\ &= Var(y_p) + E(y_p)^2 - (\delta\beta)^2 \\ &= \beta^T\Sigma\beta \end{aligned}$$

and thus

$$\begin{aligned}\rho &= \frac{\beta^T\Sigma\beta}{(\beta^T\Sigma\beta)^{1/2}(\beta^T\Sigma\beta+\sigma^2)^{1/2}} \\ &= \frac{(\beta^T\Sigma\beta)^{1/2}}{(\beta^T\Sigma\beta+\sigma^2)^{1/2}}\end{aligned}$$

We want to show that $E(E(y|y_p, X)|X) = E(E(y|y_p)|X)$. First, based on what we assume that both $y$ and $y_p$ capture all signals of $X$, we see

$$E(y|y_p, X) = E(y|X) = X\beta$$

so, the left side of equation

$$E(E(y|y_p, X)|X) = E(X\beta|X) = X\beta$$

Second, based on the bivariate normal distribution of $y$ and $y_p$ provide above, we have

$$\begin{aligned}E(y|y_p) &= \delta^T\beta + \rho\frac{(\beta^T\Sigma\beta+\sigma^2)^{1/2}}{(\beta^T\Sigma\beta)^{1/2}}(y_p - \delta^T\beta) \\ &= \delta^T\beta + \frac{(\beta^T\Sigma\beta)^{1/2}}{(\beta^T\Sigma\beta+\sigma^2)^{1/2}}\frac{(\beta^T\Sigma\beta+\sigma^2)^{1/2}}{(\beta^T\Sigma\beta)^{1/2}}(y_p - \delta^T\beta) \\ &= \delta^T\beta + (y_p - \delta^T\beta) \\ &= y_p\end{aligned}$$

so, the right side of equation

$$E(E(y|y_p)|X) = E(y_p|X) = X\beta$$

This demonstrates that both expectations are equal so:

$$E(E(y|y_p, X)|X) = E(E(y|y_p)|X) = X\beta$$

We also want to show that under the extreme case scenario, we have $Var(E(y|y_p, X)|X) = Var(E(y|y_p)|X)$ and $E(Var(y|y_p, X)|X) = E(Var(y|y_p)|X)$. First, we show $Var(E(y|y_p, X)|X) = Var(E(y|y_p)|X)$ holds true using the expectations $E(y|y_p, X)$ and $E(y|y_p)$ calculated above. On the left side of equation, we see

$$Var(E(y|y_p, X)|X) = Var(X\beta|X) = 0$$

On the right side of equation, we see

$$Var(E(y|y_p)|X) = Var(y_p|X) = 0$$

Thus, we have

$$Var(E(y|y_p, X)|X) = Var(E(y|y_p)|X) = 0$$

Second, to show $E(Var(y|y_p, X)|X) = E(Var(y|y_p)|X)$, we can show $Var(y|y_p, X) = Var(y|y_p)$. On the left side of equation,

$$Var(y|y_p, X) = Var(y|X) = \sigma^2$$

On the right side of the equation, we use the variance from the bivariate normal distribution of $y$ and $y_p$,

$$\begin{aligned} Var(y|y_p) &= (1 - \rho^2)(\beta^T \Sigma \beta + \sigma^2) \\ &= (1 - \frac{\beta^T \Sigma \beta}{\beta^T \Sigma \beta + \sigma^2})(\beta^T \Sigma \beta + \sigma^2) \\ &= \sigma^2 \end{aligned}$$

Thus, we have

$$Var(y|y_p, X) = Var(y|y_p) = \sigma^2$$

and

$$E(Var(y|y_p, X)|X) = E(Var(y|y_p)) = E(\sigma^2|X) = \sigma^2$$

In general we do not expect the machine learning prediction $\hat{f}(x)$ to perfectly estimate the entire signal in $y$, however, this extreme case serves to illustrate that the more the machine learning prediction captures the signal in the data, the closer our approximation will be. This is substantiated both by the observation that a variety of different machine learning models appear to make predictions that follow the relationship model relatively closely (see main text Figure 2) and our simulations, which show improved performance of our analytical derivation approximation as the accuracy of the machine learning model increases (see supplement Figure 1 and Figure 2).

#### 1.2 Test statistic

In section 1.1.1 we estimate the corrected coefficient $\hat{\beta}^{\star}_{(val)} = (X^T_{(val)} X_{(val)})^{-1} X^T_{(val)} (\hat{\gamma}_{0(te)} + \hat{\gamma}_{1(te)} X_{(val)} \hat{\beta}_{p(val)})$ and in section 1.1.3 we estimate the corrected conditional variance $Var(y_{(val)}|X_{(val)}) = \sigma^2_{r(te)} + \gamma^2_{1(te)} \sigma^2_{p(val)}$. The corrected coefficient and conditional variance improve from the no correction values calculated directly in the linear inferential model using the predicted outcome $y_{p(val)}$, and they are more close to the gold standard coefficient and conditional variance as if we are using the real outcomes $y_{(val)}$ in the inference model.

To form a test statistic for the corrected coefficient $\hat{\beta}_{(val)}$, we also need the standard error of $\hat{\beta}_{(val)}$. If real outcome $y_{(val)}$ are observed in the validation set and we use $y_{(val)}$ to fit a linear regression model as the inference model, we have

$$Var(\hat{\beta}_{(val)}|X_{(val)}) = (X_{(val)}^T X_{(val)})^{-1} Var(y_{(val)}|X_{(val)}) \tag{22}$$

Because in real settings the real outcome $y_{(val)}$ are unobserved, we use $\hat{\sigma}^2_{r(te)} + \hat{\gamma}^2_{1(te)}\hat{\sigma}^2_{p(val)}$ to approximate $Var(y_{(val)}|X_{(val)})$ (see details in section 1.1.3), and then $Var(\hat{\beta}_{(val)}|X_{(val)})$ can be approximated as

$$Var(\hat{\beta}_{(val)}|X_{(val)}) \approx (X_{(val)}^T X_{(val)})^{-1}(\hat{\sigma}^2_{r(te)} + \hat{\gamma}^2_{1(te)}\hat{\sigma}^2_{p(val)}) \tag{23}$$

and then,

$$se(\hat{\beta}_{(val)}|X_{(val)}) \approx \sqrt{(X_{(val)}^T X_{(val)})^{-1}(\hat{\sigma}^2_{r(te)} + \hat{\gamma}^2_{1(te)}\hat{\sigma}^2_{p(val)})} \tag{24}$$

Using the estimated corrected coefficient $\hat{\beta}^{\star}_{(val)}$ and the estimated standard error $se(\hat{\beta}_{(val)}|X_{(val)})$, we now are able to estimate a test statistic to recover the inference we would have made in the regression model Equation 1, while we actually fit the model Equation 2. To test for the null hypothesis against the alternative of the form: $H_0 : \beta_{(val)k} = 0$ vs. $H_a : \beta_{(val)k} \neq 0$ ($\beta_{(val)k}$ is the $k$-th component of the estimator vector), we can then estimate the test statistic as

$$t(\hat{\beta}_{(val)}) \approx \frac{\hat{\beta}^{\star}_{(val)}}{\sqrt{(X_{(val)}^T X_{(val)})^{-1}(\hat{\sigma}^2_{r(te)} + \hat{\gamma}^2_{1(te)}\hat{\sigma}^2_{p(val)})}} \tag{25}$$

We define a decision rule to decide whether the null hypothesis shall be rejected or not. One way is to compare the test statistic. We reject the null hypothesis $H_0 : \beta_{(val)k} = 0$ in favor of the alternative hypothesis $H_a : \beta_{(val)k} \neq 0$ at the significance level $\alpha$ when $t(\hat{\beta}_{(val)k}) > t^{\alpha}_{n-p}$ , where $t^{\alpha}_{n-p}$ is from the t statistical table with p degrees of freedom and significance level $\alpha$.

### 2 Simulation

#### 2.1 Simulation with increasing correlation between predicted and observed outcomes

In this section, we show a comparison of no correction, analytical derivation postpi, parametric bootstrap postpi and non-paramethod bootstrap postpi methods in the simulated data where the correlations between predicted and observed outcomes are different.

First, we simulate continuous covariates $x_{ij}$ and error terms $e_{u_i}$ from normal distributions, and simulate the observed outcome $y_i$ using a nonlinear function - a combination of smoothed [1] terms, quadratic and cubic terms. The model specification is:

$$\begin{aligned} x_{i1}, x_{i2}, x_{i3} &\sim \mathcal{N}(1,1) \\ e_{u_i} &\sim \mathcal{N}(0,1) \\ y_i &= \beta_1 x_{i1} + \beta_2 x_{i2}^2 + \beta_3 \cdot smooth\left(x_{i3}\right)^3 + smooth\left(e_{u_i}^2\right) \end{aligned}$$

In each simulation cycle, we set the total sample size $n = 900$, and then create a training, testing and validation set by randomly sampling the observed data into three equal size groups each with sample size 300.

On the training set, we fit a generalized additive model (GAM) [2] to estimate the prediction function $f(\cdot)$ and we use all of the covariates $x_{i1}, x_{i2}, x_{i3}$ as features to predict the observed outcomes $y_i$. On the testing set, we apply the trained prediction model to get predicted outcomes $y_{pi}$. We estimate the relationship between the observed and predicted outcome ($y_i$ and $y_{pi}$) as a simple linear regression model: $y_i \sim N(\gamma_0 + \gamma_1 y_{pi}, \sigma_r^2)$. On the validation set, we will use a linear inference model to quantify this relationship between predicted outcome $y_{pi}$ and covariate of interest $x_{1i}$.

Across 300 simulated cases, we fix the values of $\beta_2 = 1.5, \beta_3 = 1.5$ and set the standard error of the error term $e_{TSi}$ to be a range of values in $[0, 0.5, 1, ..., 6.5, 7]$. By changing this value, we are able to control the correlation between observed and predicted outcome in the testing set, in order to evaluate the performance of different methods. In the following two simulation cases, we set $\beta_1 = 0$ and $\beta_1 = 2$. In each case, we compute the estimates, standard errors, and t-statistics for $\beta_1$ with no correction, analytical derivation postpi, parametric and non-parametric bootstrap approaches. Then we evaluate the performance of above three different methods by comparing the results to the baseline results where the outcome is observed.

In the first simulation example in Figure 1, we set $\beta_1 = 0$ and compare the performance of the no correction and three postpi methods to the truth (baseline results where the outcome is observed on the validation set) under the correlation between predicted and observed outcome to be 0.1 - 0.2 to 0.7 - 0.8. The prediction has relatively little bias, so the estimated coefficients using the predicted outcome are relatively close to the estimates using the observed outcome in Figure 1(a) no matter the correlation between $y$ and $y_p$ is small or large. However, the standard errors for the no correction approach (orange color) in Figure 1(b) is much lower than what we would have observed in the observed outcomes. This is because the prediction function attempts to capture the mean function, but not the variance in the observed outcome. We show that the derivation postpi (green color), parametric bootstrap postpi (dark blue color) and non-parametric bootstrap postpi (light blue color) have values of standard error improve significantly compared to the no correction (orange color). Although the correlation between $y$ and $y_p$ is relatively small like 0.1 - 0.2, it is still obvious that the three postpi methods give the most closed standard error values compared to the truth (grey color). Because estimates are close in different methods and centered at 0 in this example, it also leads to t-statistics centered at 0 in Figure 1(c). In Figure 1(d), we show that the three postpi methods substantially correct p-values, so the distribution of p-values of our postpi methods (green color, dark blue color, and light blue color) are very similar to the distribution of the truth (grey color), but the no correction method (orange color) has skewed distribution.

In the second simulation example in Figure 2, we set $\beta_1 = 2$ and evalute the performance for different methods descrived above. In Figure 2(a), we show that the analytical derivation postpi (green color), parametric bootstrap postpi (dark blue color) and non-parametric bootstrap postpi (light blue color) provide more accurate estimate values as the correlation between predicted and observed outcome increases. We also show that the three postpi methods significantly correct standard errors in Figure 2(b) and test statistics in Figure 2(c) compared to the no correction (orange color). The corrected standard errors and t-statistics are more close to the truth (grey color) no matter the correlation between $y$ and $y_p$ is small or large. In Figure 2(d), we show the distribution of $-log10$ scale of p-values. It is obvious that the distribution of no correction (orange color) is far from the the truth (grey color), but with corrections all three postpi methods (green color, dark blue color, and light blue color) have p-value distributions more close to the truth.

### 3 Applications

#### 3.1 Predicting RNA quality

We consider another problem from the "Recount2" Project (`https://jhubiostatistics.shinyapps.io/recount/`) [3]. In this example, the phenotype we care about is RNA integrity numbers (RINs). RIN is a metric to access RNA quality, a measure of how much the RNA molecules being measured have been degraded before sequencing [4]. RIN has ranged from 1 to 10, with 1 being the most degraded RNA and 10 being the most intact [4]. RNA-seq is a powerful technique for measuring gene expression levels in cells and tissues, but it strongly relies on the quality of input RNA [5].

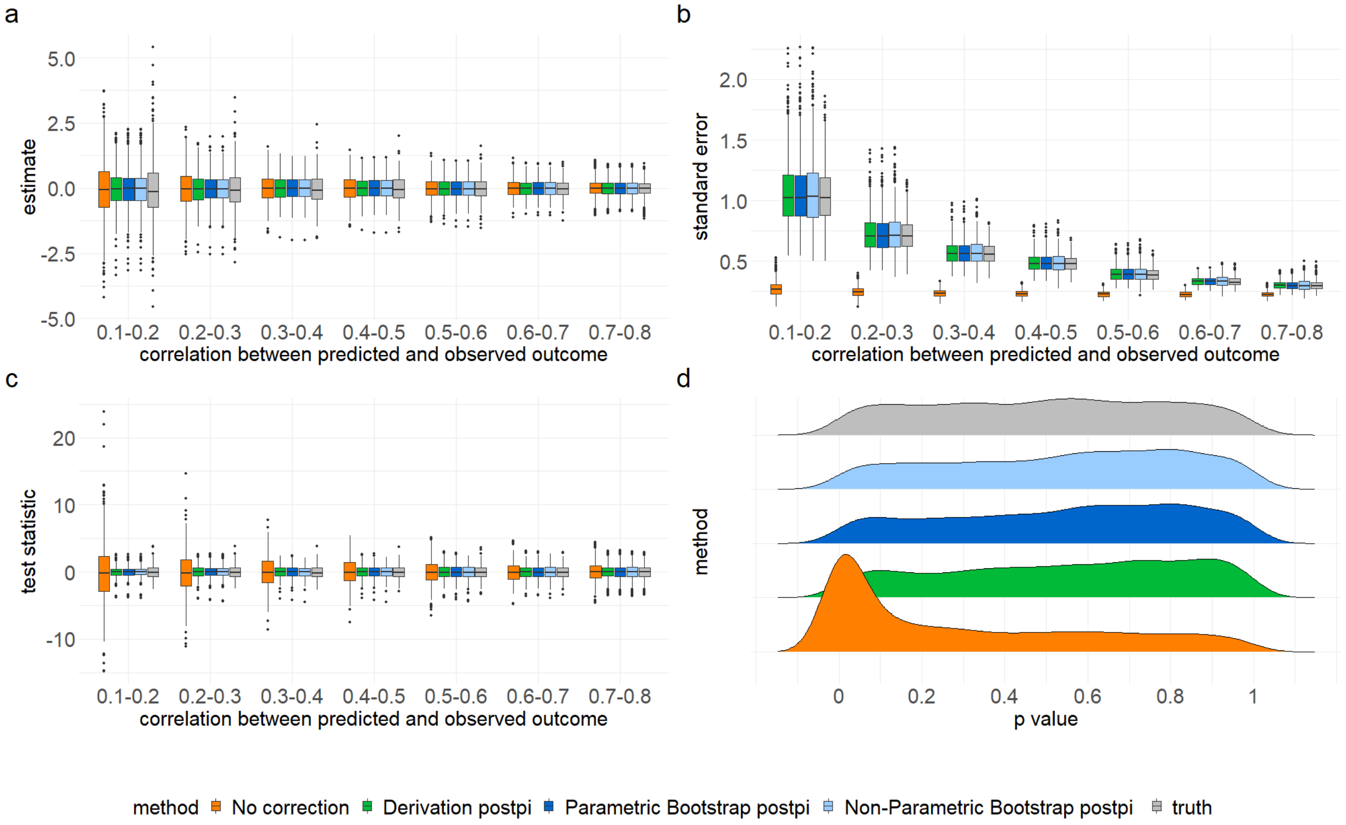


Figure 1: **Methods comparison for $\beta_1 = 0$.** Data were simulated as described in Section 2.1. On the x-axis for panel (a), (b), (c) are correlations between predicted and observed outcomes, and for panel (d) are p-values. On the y-axis we show (a) the estimates, (b) the standard errors, (c) the t-statistics, and (d) density distribution for different methods - no correction (orange color), analytical derivation postpi (green color), parametric bootstrap postpi (dark blue color), non-parametric bootstrap postpi (light blue color), and truth (grey color). The analytical derivation and bootstrap postpi approaches clearly improve the standard error and t-statistics values, and also correct the p-value distributions compared to no correction.

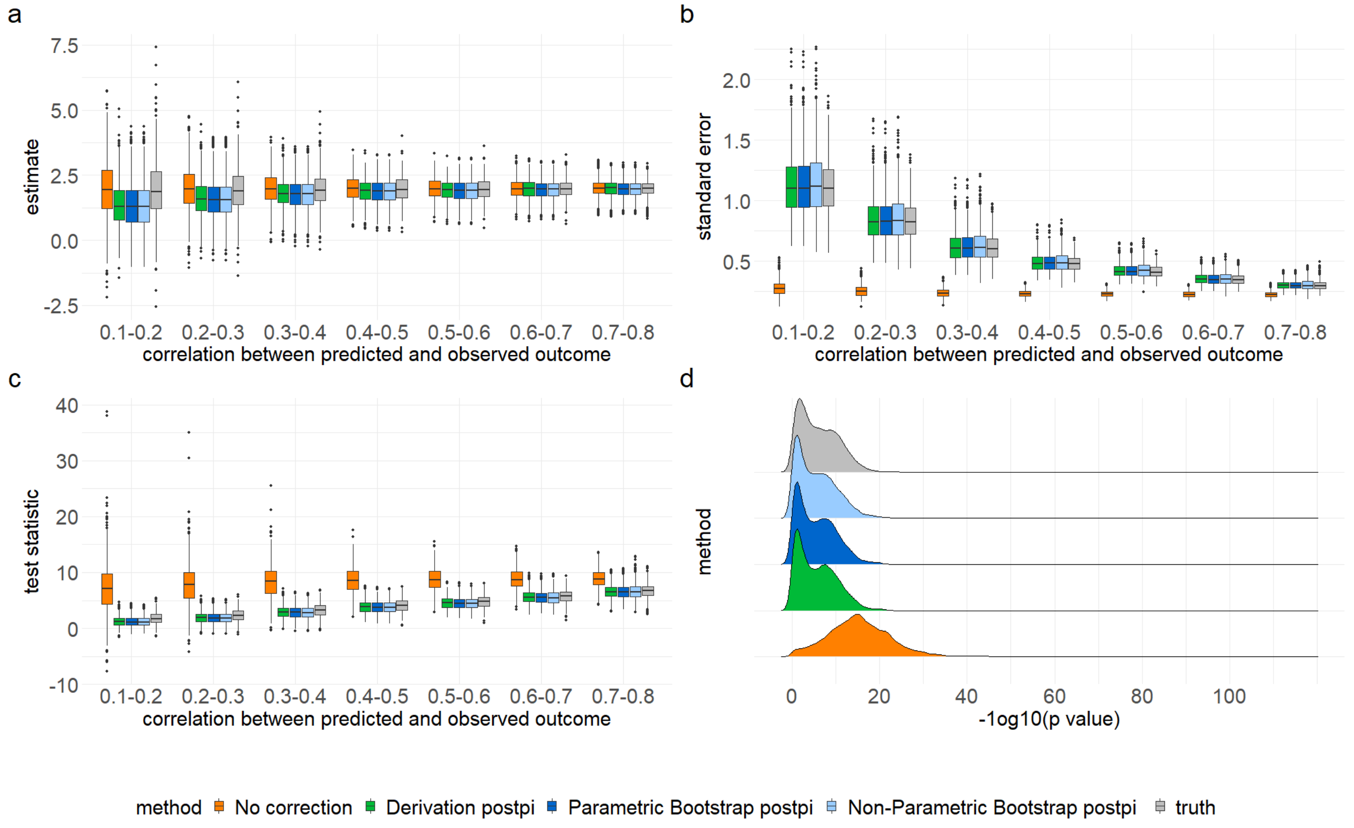


Figure 2: **Methods comparison for $\beta_1 = 2$.** Data were simulated as described in Section 2.1. On the x-axis for panel (a), (b), (c) are correlations between predicted and observed outcomes, and for panel (d) are p-values. On the y-axis we show (a) the estimates, (b) the standard errors, (c) the t-statistics, and (d) density distribution for different methods - no correction (orange color), analytical derivation postpi (green color), parametric bootstrap postpi (dark blue color), nonparametric bootstrap postpi (light blue color), and truth (grey color). The analytical derivation and bootstrap postpi approaches clearly improve the standard error values and correct the p-value distributions compared to no correction.

Because of the widespread effects of RNA quality on measurements of gene expression levels, we care about the association between RINs and gene expression levels. Studying such association is a critical step to estimate the confounding effect of RNA quality in differential expression analysis [6]. Therefore, RIN is an important technological covariate in the analysis of RNA-seq [4].

Although we have gene expression level data for all "Recount2" samples, we only observe RIN values for a small subset. However, our goal is to understand which gene expressed regions are most associated with RINs in new samples (i.e. samples without observed RINs) so that we can understand which measured genes are most impacted by RNA-quality. In this example, we collected 4769 samples from the "Recount2" where we had observed RINs, as well as the predicted RINs which were calculated from a previously trained data set using the selected 200 expressed regions as predictors [7]. In Figure 3 we show that the continuous outcomes - observed and predicted RIN values can be modeled as a simple linear relationship.

Because in a previous paper [7] we calculated the predicted values in a separate training set, in this example we only separate the dataset into a testing set with a sample size $n_1 = 2383$ and a validation set with a sample size $n_2 = 2384$. The inference model we are interested in is: $E[RIN_i|ER_i^j] = \beta_0^j + \beta_1^j ER_j$. In this model, we have $j = 1, \ldots, 200$ (expressed regions) and $i = 1, \ldots, n$, $n$ is the total number of samples in the "Recount2". Here $RIN_i$ is the RNA quality of the $i$th sample, and $ER_i^j$ is the gene expression level for the $j$th region on the $i$th sample.

On the testing set, we estimate the relationship between the observed and predicted RIN outcomes ($RIN_i$ and $RIN_{pi}$) as a linear regression model. On the validation set, we again use linear regressions as the subsequent inferential models. We fit a linear regression model to each of the available expressed regions to get 200 estimates, standard errors, and t-statistics. We then compare the analytical derivation postpi, parametric bootstrap postpi, nonparametric bootstrap postpi, and the no correction approaches we did with the simulated data.

We observed that the estimates are quite similar among the three approaches in Figure 4(a) where RMSE for no correction (orange color) is 0.012 compared to the truth, 0.019 for analytical derivation postpi (green color), 0.018 for parametric (dark blue color) and non-parametric (light blue color) bootstrap postpi methods. The standard errors were underestimated by no correction (orange color) in Figure 4(b) with RMSE 0.0015, improved to 0.0014 for non-parametric bootstrap postpi (light blue color), and further reduced to 0.00008 for analytical derivation postpi (green color) and 0.00009 for parametric bootstrap postpi (dark blue color). The resulting t-statistics are in Figure 4(c) where RMSE for no correction (orange color) is 1.71 compared to the truth and

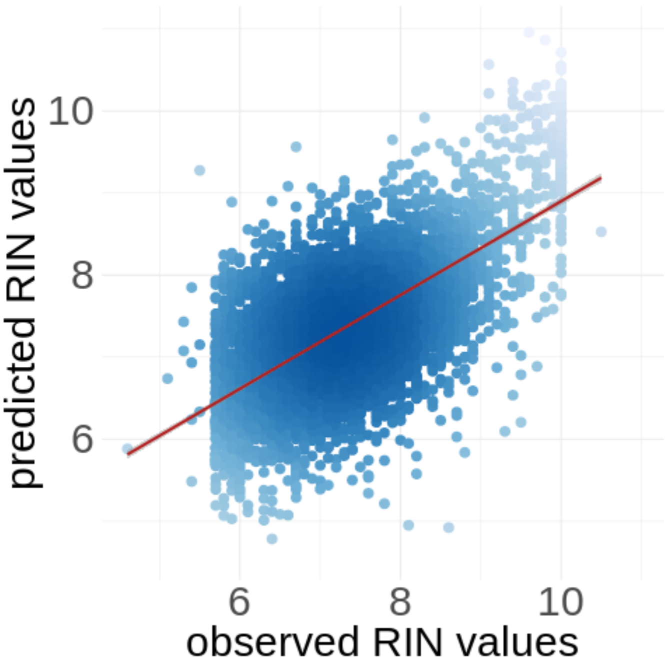


Figure 3: **Relationship between the observed and predicted RNA Quality.** Data were collected from the "Recount2" Project as described in Section 3.1. On the x-axis are the observed RIN values and on the y-axis are the predicted RIN values on the validation set. We observe that the observed and predicted RIN values can be modeled as a simple linear model.

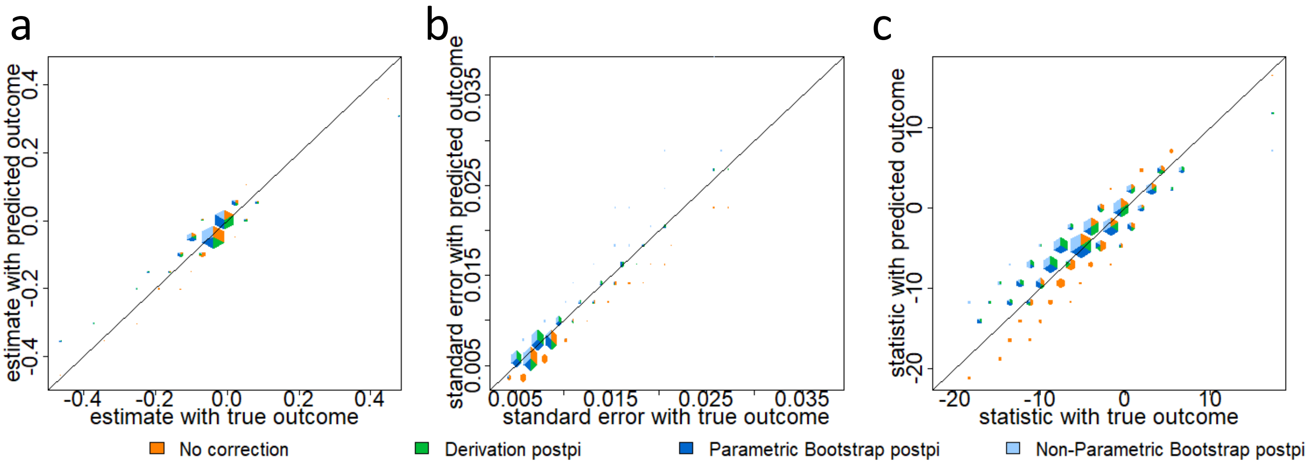


Figure 4: **RNA quality prediction.** Data were collected from the "Recount2" Project as described in section 3.2.1. On the x-axis are the values calculated using the observed outcome and on the y-axis are the values calculated using no correction (orange color), analytical derivation postpi (green color), parametric bootstrap postpi (dark blue color), and non-parametric bootstrap postpi (light blue color). We show (a) the estimates are similar across all four approaches since the data were simulated from a normal model, (b) the standard errors are small for the uncorrected inference (orange color) but corrected with our approaches and (c) the t-statistics are anti-conservatively biased for uncorrected inference but corrected with our approaches.

improved to 1.66 for postpi bootstrap parametric (dark blue color). The analytical derivation postpi (green color) methods has RMSE 1.83 and non-parametric bootstrap postpi (light blue color) has RMSE 1.95. The analytical derivation postpi, parametric and non-parametric bootstrap postpi methods attenuate the signal when the estimates are extreme but is accurate when the estimates should be small, and all three postpi methods accurately correct the standard error of estimates. This is not surprising because we are introducing variability from the prediction model into the estimates.

### References

[1] R Core Team. *R: A Language and Environment for Statistical Computing.* R Foundation for Statistical Computing, Vienna, Austria, 2018.

[2] Simon N Wood. Fast stable direct fitting and smoothness selection for generalized additive

models. *Journal of the Royal Statistical Society: Series B (Statistical Methodology)*, 70(3):495–518, 2008.
